# On the hormonal control of posterior regeneration in the annelid *Platynereis dumerilii*

**DOI:** 10.1101/2022.01.21.477205

**Authors:** Patricia Álvarez-Campos, Anabelle Planques, Loïc Bideau, Michel Vervoort, Eve Gazave

**Affiliations:** Université de Paris, CNRS, Institut Jacques Monod, F-75013, Paris, France; Centro de Investigación en Biodiversidad y Cambio Global (CIBC-UAM) & Departamento de Biología (Zoología), Facultad de Ciencias, Universidad Autónoma de Madrid, Spain

**Keywords:** regeneration, brain hormone, *Platynereis*, annelid, methylfarnesoate, hormonal regulation

## Abstract

Regeneration is the process by which many animals are able to restore lost or injured body parts. After amputation of the posterior part of its body, the annelid *Platynereis dumerilii* is able to regenerate the pygidium, the posteriormost part of its body that bears the anus, and a subterminal growth zone containing stem cells that allows the subsequent addition of new segments. The ability to regenerate their posterior part (posterior regeneration) is promoted in juvenile worms by a hormone produced by the brain and is lost when this hormonal activity becomes low at the time the worms undergo their sexual maturation. By characterizing posterior regeneration at the morphological and molecular levels in worms that have been decapitated, we show that the presence of the head is essential for multiple aspects of posterior regeneration, as well as for the subsequent production of new segments. We also show that methylfarnesoate, the molecule proposed to be the brain hormone, can partially rescue the posterior regeneration defects observed in decapitated worms. Our results are therefore consistent with a key role of brain hormonal activity in the control of regeneration and growth in *P. dumerilii*, and support the hypothesis of the involvement of methylfarnesoate in this control.

## Introduction

Regeneration, the ability to restore lost or injured body parts, re-establishing both morphology and function (Brockes & Kumar 2008; Poss 2010), is a widespread phenomenon in Metazoa (Bely & Nyberg 2010; Bideau et al. 2021). However, regeneration in animals is also highly variable as some species are able to regenerate only specific cell types or tissues, while others can regenerate complex structures such as limbs, and even their whole body from a small piece of tissue (Bely & Nyberg 2010; Grillo et al. 2016; Bideau et al. 2021). Segmented worms (Annelida) have been widely used to study this amazing process and its evolution in animals (*e*.*g*., Zattara & Bely 2016, Özpolat & Bely 2016). Many annelids (with the noticeable exception of leeches) display substantial regeneration abilities, which however may differ from one species to another. In fact, even in closely-related species, a huge variability is observed, in particular with respect to the possibility of regenerating their anterior body region upon amputation (Özpolat & Bely 2016, Nikanorova et al. 2020). Moreover, regeneration in some semelparous annelids (*i*.*e*., species with a single reproduction episode followed by death), such as nereids (Nereidae), is intimately linked to its sexual reproduction mode, given that these species lose their growing and regenerative abilities when sexual maturation starts (*e*.*g*., Clark & Ruston 1963; Clark & Scully 1964; Schenk et al. 2016).

Nereids, such as *Hediste diversicolor* (formerly *Nereis diversicolor*), *Alitta virens* (formely *Nereis virens*) and *Platynereis dumerilii*, are indeed able to grow during most of their life (juvenile phase), by adding segments in their posterior body region one by one, a process known as posterior growth or posterior elongation (*e*.*g*. Clark & Scully 1964; de Rosa et al. 2005; Gazave et al. 2013). During this growth phase, these worms are also able to successfully regenerate the posterior part of their body upon amputation, a process known as posterior regeneration (*e*.*g*., Clark & Ruston 1963; Özpolat & Bely 2016; Kozin & Kostyuchenko 2015; Planques et al. 2019). In contrast, anterior regeneration in nereids is quite limited or unsuccessful (Hauenschild 1960), only occurring if at least a small part of the prostomium (*i*.*e*., the anterior-most portion of the body that comprises a specific part of the brain, the supraoesophageal ganglion) is still present (Hofmann 1975). Interestingly, both posterior growth and posterior regeneration abilities stop when the worms undergo sexual maturation, a process known as epitoky, which involves dramatic modifications of the morphology, physiology, and behaviour of the worms to become reproductive gonochoristic individuals (*e*.*g*. Chatelain et al. 2008). During this sexual metamorphosis, the benthic tubicolous juvenile worms turn into pelagic reproductive individuals, undergoing striking modifications, such as histolysis of muscles, gut degeneration, enlargement of appendages (parapodia), formation of special natatory chaetae, acquisition of sex-specific body coloration (yellow females and red-whitish males, thanks to respective gametes colours), and production of massive amounts of gametes (*e*.*g*., Fauvel 1959; Clark 1961; Fischer 1999). Sexually mature animals are then ready to spawn, and they start a nuptial dance (females and males rapidly swim together in a circle) to deliver their gametes and die shortly after spawning.

A large series of experiments have shown that sexual maturation in nereids is under an endocrine control and identified the supraesophageal ganglion as the source of this hormonal activity (*e*.*g*., Durchon 1948; Hauenschild 1964; Bertout 1983; Fischer 1984; Schenk et al. 2016). In juvenile worms, the brain hormone, traditionally termed nereidin, was shown to repress sexual metamorphosis. Indeed, sexual maturation is triggered by the decrease of the effective concentration of the brain hormone, which occurs when the worms become older and during the breeding season (*e*.*g*., Clark & Ruston 1963; Scully 1964; Golding 1983). In addition to repressing epitoky, the brain hormone was also suggested to promote both posterior growth and posterior regeneration. Indeed, if the supraoesophageal ganglion is totally removed from nereid worms before posterior amputation, posterior regeneration and subsequent posterior growth are impaired (*e*.*g*., Clark & Bonney 1960; Clark & Evans 1961; Clark & Ruston 1963; Durchon & Marcel 1962; Hauenschild 1966; Golding 1967a; Hofman 1976). Remarkably, these effects can be reversed by implanting supraoesophageal ganglia from intact worms in decerebrated hosts, which implies that the hormone stimulating regeneration and growth is continuously produced by the brain, even in the absence of any amputation (*e*.*g*. Scully 1964; Golding 1967a). The molecular identity of the brain hormone has remained elusive for many years. While initially believed to be a neuropeptide (Müller 1973; Hofmann 1976; Cardon et al. 1981; Durchon 1984), it was recently found that nereidin activity in *P. dumerilii* involves methylfarnesoate (MF), a sesquiterpenoid hormone related to arthropod juvenile hormones (Schenk et al. 2016). MF was indeed shown to repress vitellogenesis in coelomic cells, which was used as a read-out of sexual maturation, and its concentration drops in the course of maturation, as expected for nereidin (Schenk et al. 2016). MF treatment also increased the expression of *Hox3*, a gene expressed in the growth zone in both growing and regenerating *P. dumerilii* worms (Pfeifer et al. 2012; Gazave et al. 2013; Planques et al. 2019), which led to the hypothesis that MF might also promote posterior growth and regeneration (Schenk et al. 2016).

In this article, we further studied the hormonal control of posterior regeneration in *P. dumerilii*. In recent years this small-sized nereid annelid has become an important model for comparative neurobiology and developmental biology (*e*.*g*., Williams and Jékely 2016; Schenkelaars & Gazave 2021; Özpolat et al. 2021). In three days, embryonic and larval development gives rise to small juvenile worms that display a head, three segments with a pair of parapodia and a terminal body element called the pygidium, which bears the anus and characteristic paired outgrowths (anal cirri) (Fischer et al. 2010). These worms subsequently enter a posterior growth phase that spans over from five days to more than one year (in our laboratory culture conditions). This juvenile posterior growth phase ends when the worms metamorphose into sexually mature individuals (from four months to more than one year old) (Fig. 1). Posterior growth relies on the presence and activity of a subterminal growth zone, located immediately in front of the pygidium, which contains putative stem cells whose sustained divisions allow the serial addition of new segments in the posterior body region (de Rosa et al. 2005; Gazave et al. 2013).

**Figure 1.**
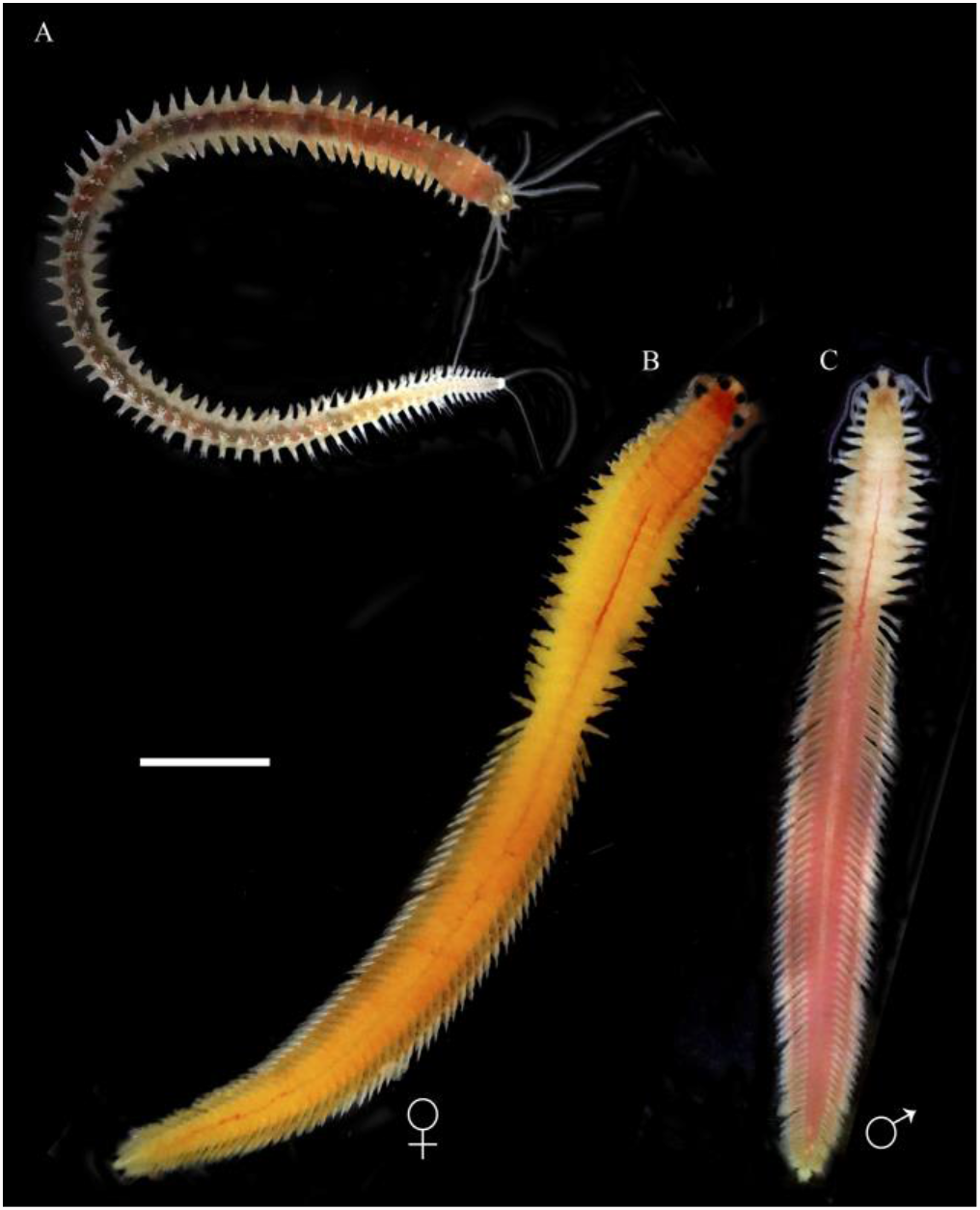
Morphology of *P. dumerilii* worms. Juvenile (A) and sexually mature (female, B and male, C) worms are shown. Scale bar = 500 μm.

During this juvenile growth phase, after a posterior amputation which consists of the removal of several segments, the growth zone, and the pygidium, *P. dumerilii* worms are able to quickly reform their pygidium and their growth zone that in turn produces new segments that replace the amputated ones. We recently showed that posterior regeneration in *P. dumerilii* is a rapid process that includes five well-defined stages in a reproducible timeline (Planques et al. 2019). As shown in Planques et al. 2019, and illustrated in supplementary Fig. 1, stage 1, which is reached one day post posterior amputation (1 dppa), corresponds to the completion of wound healing. At stage 2 (2 dppa), the anus opening is reformed and a small blastema is observed. From this stage onwards, intense cell proliferation can be detected in the regenerated region. The size of the blastema increases at stage 3 (3 dppa) and the regenerated pygidium starts to differentiate as shown by the formation of small anal cirri and pygidial muscles. The analysis of the expression of molecular markers, such *Hox3* and *engrailed*, indicates that the growth zone has already been regenerated at this stage and produced the primordium of at least one segment. The size of the regenerated region continues increasing during the two next and last steps of regeneration (stage 4 and 5; 4 and 5 dppa, respectively). At stage 4, a well-differentiated pygidium with long anal cirri is observed. At stage 5, the boundary between the pygidium and the new developing segments becomes clearly visible through the appearance of lateral segmental grooves. At this stage, posterior regeneration is finished. In the following days, the worms enter a post-regenerative posterior growth phase and can be scored by the number of segments with conspicuous boundaries and developing parapodia that are present. While the timeline of regeneration stages is highly reproducible in worms of a given age and size, the subsequent addition of segments is much more variable (Planques et al. 2019).

In this article, we took advantage of our previous morphological and molecular detailed characterizations of posterior regeneration in *P. dumerilii* to re-address the role of the brain hormonal activity in this process. By combining anterior and posterior amputations and performing both morphological and molecular characterizations on precisely staged worms, we found that both posterior regeneration and post-regenerative posterior growth are severely affected in the absence of the brain, and characterized these defects at the morphological and molecular levels. We also observed that posterior regeneration defects in decapitated worms can be partially rescued by applying exogenous MF hormone, pointing out the key role of MF in the control of posterior regeneration.

## Materials and methods

### Worm culture, amputation procedures and production of the biological material

Individuals of *P. dumerilii* were obtained from the laboratory culture established at the Institut Jacques Monod (France), following Dorresteijn et al. (1993) protocol (Vervoort & Gazave 2021). In all but one experiment, 3-4-month-old worms with 30-40 segments were used. In addition, in the sexual maturation experiment, 1-month-old worms with 10–15 segments and 1-year-old worms with 85–95 segments were also used. After anaesthesia (with a solution of MgCl_2_ 7.5% and sea water, 1/1), amputations of the anterior part of the worms were done using a microknife (SharpointTM) at the anterior boundary of the first segment in order to remove the prostomium (including the supraesophageal gland). Amputations of the posterior-most segments of 3-4-month-old worms were also performed in order to remove the last 5-6 segments, the growth zone and the pygidium (Planques et al. 2019). After amputation, all worms were let to recover in natural fresh sea water (NFSW) for a couple of minutes and were then placed individually in 6-well plates in 10 ml of NFSW.

### Methylfarnesoate treatments

Following Schenk *et al*. (2016) experimental design, (E,E)-methylfarnesoate (MF; Echelon Biosciences) 100 nM stock was prepared by adding 1 μl of pure hormone in 33.6 ml of DMSO in a glass container to prevent potential hydrophobic binding of the hormone to plastic. MF stock was then aliquoted in 10-12 glass vials that were frozen at -20ºC, using only one aliquot per day of experiment in order to avoid MF alteration after several freeze-thaw cycles. Five to six amputated worms (see above) were placed in 150 ml glass beakers. MF-treated worms were maintained in a solution of 100 ml filtered NFSW and 100 μl of MF (from 100 nM stock solution in 0.1% DMSO). Worms used as controls (non-MF-treated, with or without anterior amputation) were maintained in 100 ml filtered NFSW and 0.1% DMSO.

### Scoring of the worms and statistical analyses

To monitor the morphological changes expected for sexual maturation following anterior amputation, as well as the posterior regeneration stages following anterior and/or posterior amputations, daily observations of the worms were done after amputation and/or hormonal treatments (see the “Results” section for the different timing of scoring/imaging procedures) using a ZEISS dissecting scope (Stemi SV11). Scoring was done according to the staging system established in Planques *et al*. (2019): worms were scored either by the regeneration stage that has been reached (stage 1 to 5) or, after regeneration completion, by the number of new segments that has been produced during the post-regenerative posterior growth process. During regeneration, worms showing a morphologically intermediate stage were coded as 1.5, 2.5, 3.5, and 4.5. Worms that died in the middle of the experiment were counted but removed from the statistical analyses, and those with abnormal morphology were analysed separately, to avoid biased results. Statistical analyses and graphic representation of the results of the amputation experiments were done using Prism 7 for Mac OS (GraphPad software) (Ivashchenko et al. 2017). Detailed results of all the statistical analyses are provided in Supp. Table 1.

### *Whole-mount* in situ *hybridizations (WMISH) and imaging*

Single nitro blue tetrazolium chloride/5-bromo-4-chloro-3’-indolyphosphate (NBT/BCIP) WMISH on regeneration stages were performed as previously described (*e*.*g*., Tessmar-Raible et al. 2005; Gazave et al. 2017; Planques et al. 2019). Bright field pictures for NBT/BCIP WMISH were taken on a Leica microscope DM5000B. Editing and compilation of Z projections were achieved using ImageJ and Adobe Photoshop CS5.1. The final figure panels were then compiled using Adobe Illustrator CS5.1.

## Results

### Timeline of events following anterior amputation

In order to be able to identify the effects of anterior amputation and brain removal on *P. dumerilii* posterior regeneration, we first characterized their consequences on worm survival and sexual maturation. As a consequence of the removal of the supraesophageal ganglion that produces brain hormones preventing sexual maturation, decapitation has indeed been shown to lead to accelerated sexual maturation of *P. dumerilii* worms, which is followed by the death of the animals, (*e. g*., Hauenschild 1960, 1964, 1966). Effective concentration of brain hormones is thought to decrease from younger to older animals (*e*.*g*., Golding 1967a, 1983), raising the possibility that decapitation could have age-dependent effects (or timelines) due to different hormone concentrations in the worms’ tissues. We therefore did a series of decapitation experiments using 1-month-old (supposedly high hormone level), 3-4-month-old (supposedly intermediate hormone level), and 1-year-old worms (supposedly low hormone level). Worms were observed at many different time points after anterior amputation. For the three conditions, non-amputated worms of the same age were used as controls. As expected, while some 1-year-old control worms became sexually mature during the course of the experiment, we did not observe any signs of sexual maturation in 1-month-old and 3-4-month-old control animals (not shown).

First of all, we found that the vast majority of decapitated worms survived for several days after amputation. Death occurred from 12 or 14 days post anterior amputation (dpaa) for 1-month-old and 3-4-month-old worms, respectively (in agreement with Hauenschild & Fischer, 1962), or from 21 dpaa for 1-year-old animals (Fig. 2). Interestingly, regardless of their age, all decapitated worms showed some signs of sexual maturation after anterior amputation. From 2-3 dpaa onwards, large cells (lc) are visible near the amputation site in the three categories of worms (Fig. 2A1, B1, C1). Those cells, which might be maturing germ cells, started to accumulate in the whole body at 8-10 dpaa in 1-month-old and 3-4-month-old animals (Fig. 2A2, B2). In addition, in those worms, the accumulation of large cells coincided with the enlargement of parapodia (Fig. 2A2, A3, B2), a typical feature of sexual maturation. Probably due to the thickness of the body wall, the accumulation of large cells throughout the body could not be noticed as convincingly in the 1-year-old worms (Fig. 2C1). The enlargement of parapodia in these individuals was visible regardless from 8 dpaa onwards (Fig. 2C2). Concomitantly, and only in 1-year-old worms, the typical color differences between males and females started to appear (Fig. 2C3, C4). In addition, at 12 dpaa, 1-year-old worms showed an accumulation of large cells that we identified as gametes based on previous descriptions of *P. dumerilii* mature oocytes (see figure 5 in Hauenschild & Fischer, 1962). Indeed, those cells are very different between females (yellowish individuals) and males (reddish individuals) and are morphologically similar to oocytes and sperm observed in non-amputated worms that underwent sexual maturation (Fig. 2C5, C6). From around 19 dpaa onwards, 1-year-old decapitated animals showed the typical *P. dumerilii* courtship dancing, yet none of the worms managed to release their gametes and perform fertilization. In contrast, in 1-month-old and 3-4-month-old worms that were anteriorly amputated, we did not observe any accumulation of gametes in the worms, nor any signs of courtship behavior.

**Figure 2.**
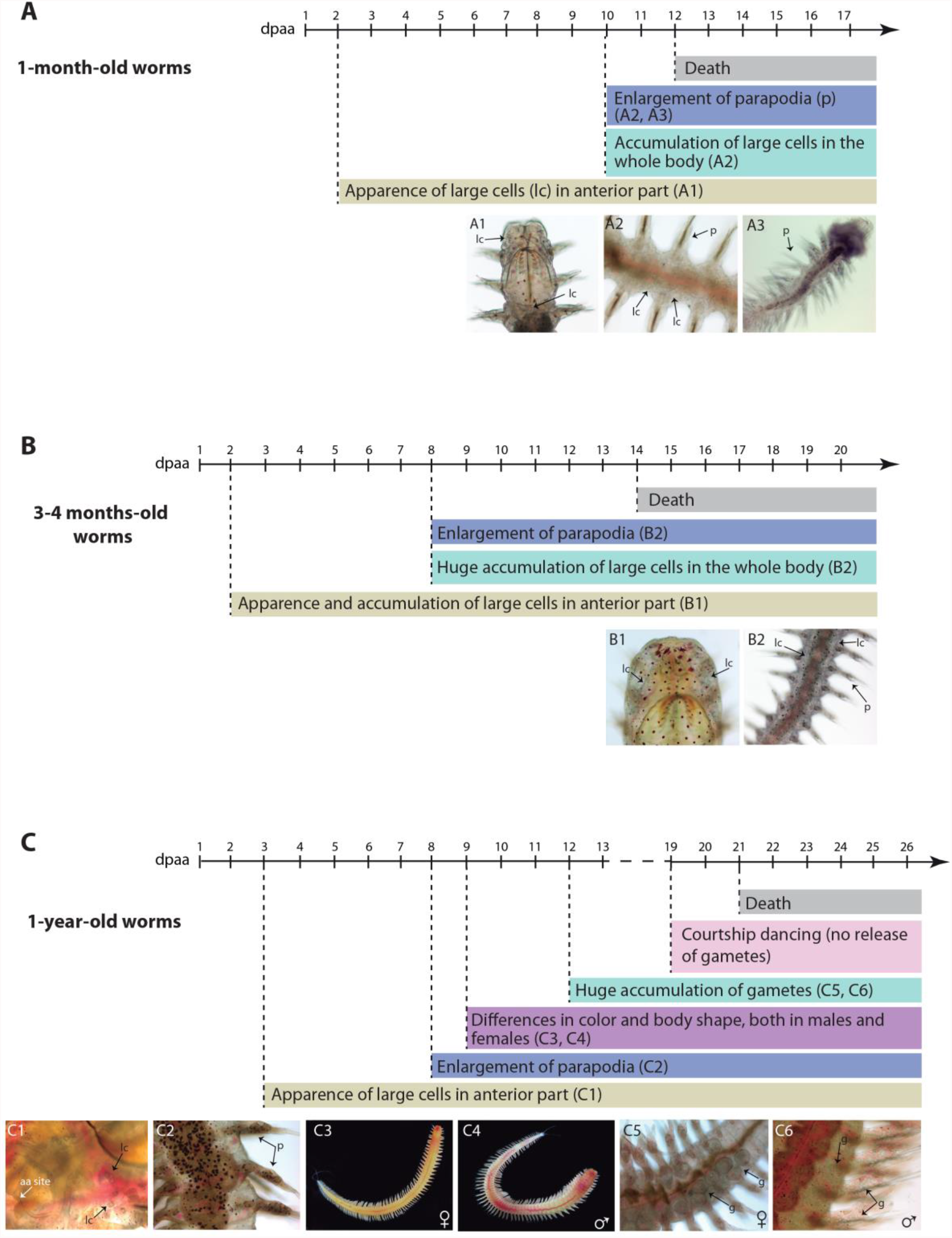
Effects of anterior amputation on *P. dumerilii* worms’ survival and sexual maturation. Three categories of worms were studied: 1-month-old (A), 3-4-month-old (B) and 1-year-old worms (C). For each panel (A to C), a timeline in days post anterior amputation (dpaa), as well as specific morphological features observed (with a representative picture) and their duration, are provided. A to C) At 2-3 dpaa, large cells (lc), that may be germ cells, started to appear near the amputation site (aa site) (A1, B1, C1). These cells accumulated in the whole body at around 8-10 dpaa, especially in 1- and 3-4-month-old worms (A2, B2). At the same time, parapodia (p) started to enlarge (A2, A3, B2, C2), and the typical color differences exhibited between mature females (yellowish) and males (pink-reddish) were patent in 1-year-old individuals (C3, C4). At 12 dpaa, 1-year-old individuals exhibit accumulation of gametes (g) (C5, C6). The death of the decapitated worms occurs from 12 to 21 dpaa, depending on their age.

In conclusion, we found that anterior amputation does indeed induce sexual maturation in 1-year-old worms and that the decapitated animals were however unable to reproduce. Nevertheless, decapitated 1-year-old individuals were able to do the courtship dancing, suggesting that the control of this behavior is not dependent on the prostomium as it was previously hypothesized (Boilly-Marer 1969a, b). In 1-month-old and 3-4-month-old worms, only some limited aspects of sexual maturation (possible accumulation of undifferentiated germ cells and parapodia enlargement) were observed (Fig. 2A, B). Importantly, decapitated worms were able to survive for more than 10 days, which made it possible to combine anterior and posterior amputations to assess the effects of the absence of the brain on posterior regeneration.

### Morphological characterization of posterior regeneration after both anterior and posterior amputations

Seminal studies performed in the fifties and sixties convincingly showed that the removal of the brain impaired posterior regeneration in nereid annelids such as *P. dumerilii* (see Introduction for details and references). However, in these studies, animals were not always precisely staged (no systematic selection of specific age or size worm to limit intra-individual variability), as taken often directly from the wild, and, importantly, no distinction was generally made between regeneration *per se* (pygidium and growth zone reformation) and post-regenerative posterior growth (segment addition from the regenerated growth zone). To reassess the influence of the absence of the brain hormone on posterior regeneration in *P. dumerilii*, we therefore decided to perform experiments combining anterior amputation followed by posterior amputation. We used 3-4-month-old worms, for which we previously established a precise timeline of regeneration stages (Planques *et al*. 2019) and that are able to survive for 14 days after anterior amputation (see previous section). As we did not know for how long significant concentrations of the brain hormone were maintained in the blood and tissues after brain removal, we performed anterior amputation (Fig. 3A, black diamond) at different time points before posterior amputation (Fig. 3A, red diamond). Seven experimental conditions (for a total of 316 amputated individuals) plus one control experiment (only posterior amputation) were designed for a total duration ranging from 7 to 14 days (Fig. 3A). For all experiments, day 0 was defined as the day when posterior amputation was performed. In condition C0, both anterior and posterior amputations were performed the same day (day 0). In condition C1, anterior amputation was performed one day before posterior amputation (day -1). Thus, in C1 there is a one-day of delay between anterior and posterior amputations. C2 to C7 experiments were similarly designed with anterior amputations performed two to seven days before posterior amputation (Fig. 3A). Posterior regeneration stages reached by the worms, or the number of segments that has been added for late time points, were scored every day for seven days post posterior amputation (dppa) (Fig. 3A). Morphology of the regenerated region was also monitored and abnormal individuals with morphologies difficult to assign to a precise stage were excluded from the main statistical analysis and analyzed separately (Fig. 4).

**Figure 3.**
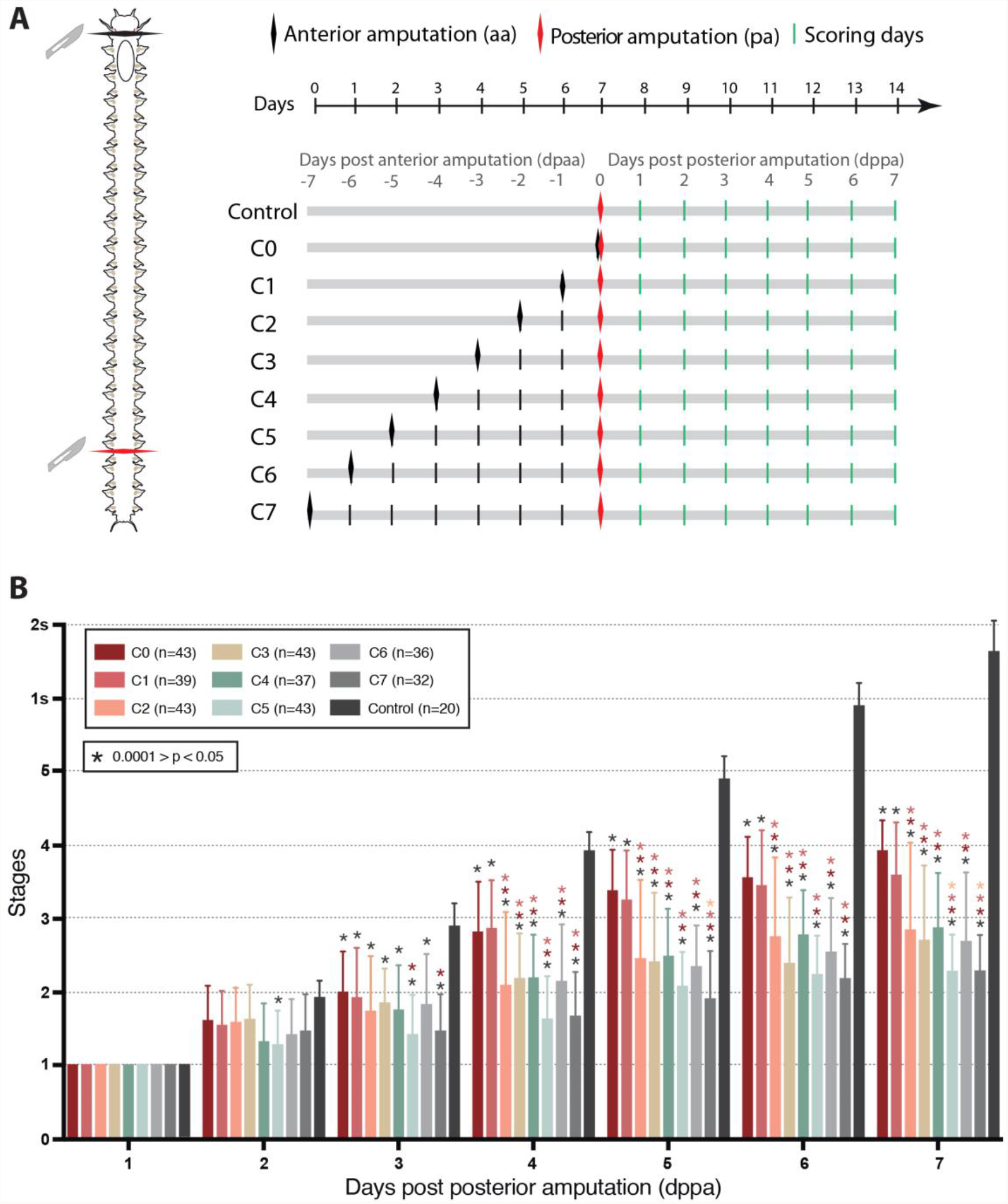
Influence of anterior amputation on *P. dumerilii* posterior regeneration. A) Schematic representation of the experimental design used to address the impact of anterior amputation (aa, black diamonds) on the regeneration that follows posterior amputation (pa, red diamonds). A timeline representing the whole duration of the experiment (14 days) and the timepoints for anterior amputation (aa) and posterior amputation (pa) for each condition, as well as the scoring days (green bars, from 1 to 7 days post posterior amputation, dppa), are provided. Control worms were only amputated posteriorly; for C0, both anterior and posterior amputations were done the same day (0 dppa); for C1 to C7, anterior amputation was made 1 to 7 days before posterior amputation (−1 to -7 dppa). B) Graphic representation of the regeneration stages reached by the worms, every day for 7 days after posterior amputation, for each condition (see inset for the color code and the number of worms used per condition). Asterisks above the bar indicate that this condition was significantly different to the other conditions specified by the asterisk color. Statistics: 2-way ANOVA on repeated measures with Tukey correction. * 0.0001 > p < 0.05 (for specific p-values, see Supp. Table 1). Error bar: Standard Deviation (SD).

**Figure 4.**
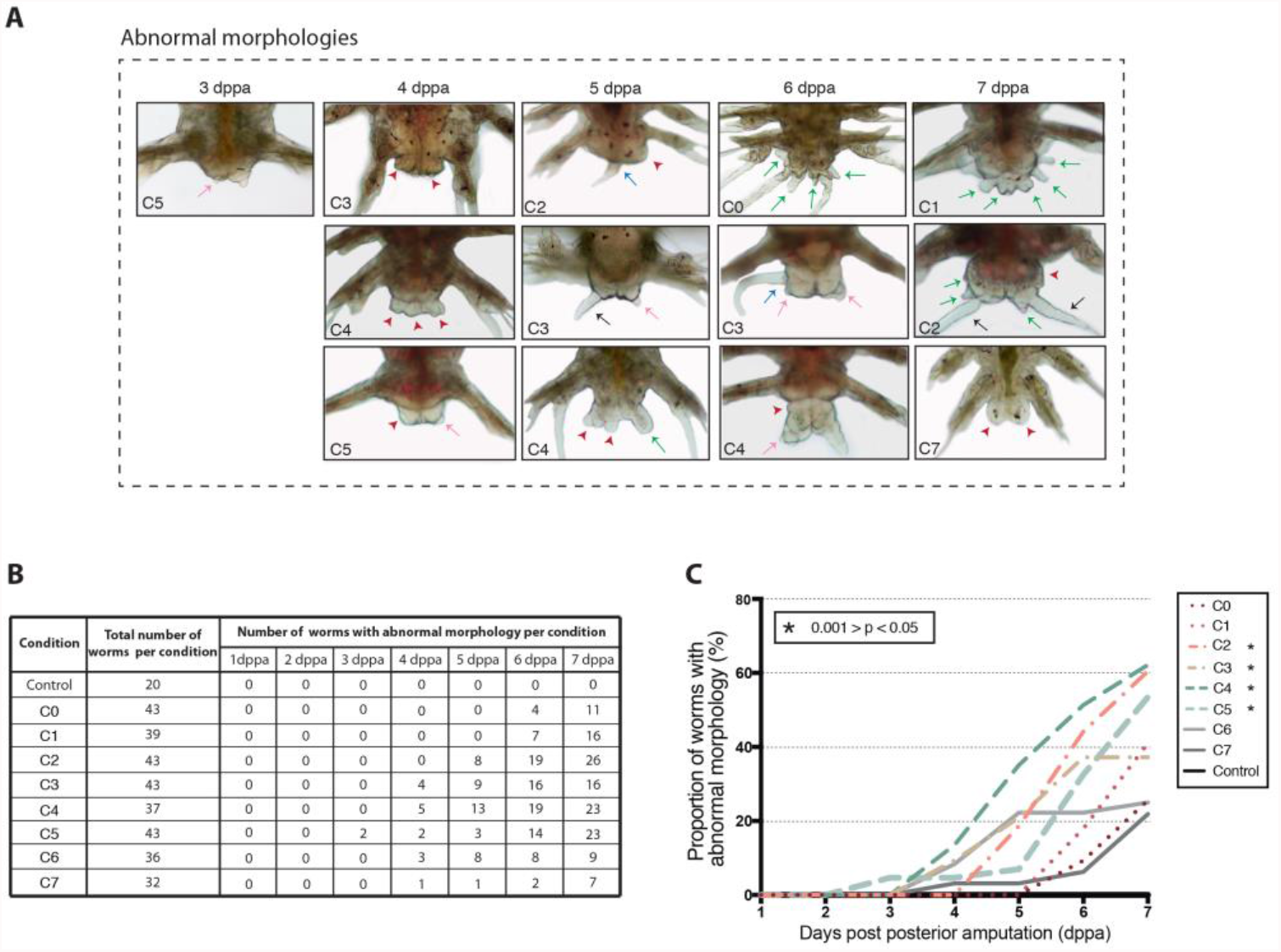
Abnormal morphologies observed during *P. dumerilii* posterior regeneration after anterior amputation. A) Bright-field microscopy images of the most striking abnormal morphologies observed on posterior regeneration at different time points (4 to 7 days post posterior amputation or dppa). For each abnormal morphology shown, the condition of anterior amputation performed (C0 to C7, see Figure 3 and text for details) are mentioned on the image. Black arrows = well-developed anal cirri; red arrowheads = enlargement of regenerated region; pink arrows = reduction of anal cirri; green arrows = additional tiny outgrowths; blue arrows = misplaced anal cirri. B) The number of worms showing abnormal morphologies during posterior regeneration in each different condition (C0-C7 and control) is provided for each scoring day (from 1 to 7 dppa). No worms with abnormal morphology were found in the control condition. The total number of worms per experimental condition is mentioned. C) Graphic representation of the proportion of worms showing abnormal morphologies on posterior regeneration for each condition of anterior amputation (C1 to C7, and control) from 1 to 7 dppa. Asterisks indicate significant differences in comparison to the control, as defined by 2-way ANOVA on repeated measures with Dunnett correction. * 0.0001 > p < 0.05 (for specific p-values, see Supp. Table 1).

The overall results of these experiments are depicted in Fig. 3B. Detailed statistics can be found in Supp. Table 1. Stage 1, which corresponds to the completion of wound healing, was reached at 1 dppa in all conditions like in the control condition. No morphological abnormalities were observed at this stage, indicating that wound healing is not affected by the absence of the head. Worms in all conditions also reached stage 2 which is characterized by the presence of a regenerated anus and a small blastema. Worms from all but one experimental condition started to be significantly delayed in their posterior regeneration at 3 dppa (C5 worms were already significantly different from controls at 2 dppa) and none of these worms were able to complete posterior regeneration (*i*.*e*., to reach stage 5) and produce segments at the end of the experiment (at 7 dppa), in contrast to the controls. The importance of the posterior regeneration delay was correlated to the experimental condition, *i*.*e*. the moment when anterior amputation is performed. Indeed, C0 and C1 worms were the less affected ones and they were also significantly different from all the other conditions from 4 dppa onwards (Fig. 3B, Supp. Table 1). Some C0 and C1 worms reached stage 4 at 7 dppa, unlike worms from all the other experimental conditions. The most affected conditions were C5 and C7 which were significantly delayed at 7 dppa as compared to C0, C1, and C2.

In addition to a delay in posterior regeneration, various abnormal morphologies not observed in the control worms were also observed in all experimental conditions from 3 dppa onwards (Fig. 4). The most affected structures were the anal cirri (Fig. 4A, black arrows). From 3 dppa onwards, several worms displayed an enlarged regenerated region (Fig. 4A, red arrowheads) and some showed one or two outgrowths at the posterior end of the pygidium, that were reduced in size (Fig. 4A, pink arrows) or misplaced (Fig. 4A, blue arrows). In some worms, additional tiny outgrowths often located next to two well-developed anal cirri, were also observed (Fig. 4A, green arrows). C5 was the only condition in which worms with abnormal morphologies were observed at 3 dppa (Fig. 4B), in a proportion of ∼5% of the total number of analyzed worms (Fig. 4C). At 4 dppa, worms with abnormal morphologies were found in C3 to C7 conditions (Fig. 4B, C), with a proportion ranging from ∼3% (in C7) to ∼13% (in C4) of the total number of analyzed worms (Fig. 4B, C). Abnormalities were also observed in C2 at 5 dppa (∼18%), and in C0 and C1 at 6 dppa (∼9% and ∼18%, respectively) (Fig. 4B, C). At the end of the experiment (7 dppa), the conditions that resulted in the highest number of abnormal regenerated worms were C2, C4, and C5, with 26, 23, and 23 worms, respectively (Fig. 4B), which corresponds to ∼60% of the total observed specimens (Fig. 4C).

In conclusion, our data show that, while not affecting early steps of regeneration, the absence of the head prevents regeneration from proceeding beyond stage 3 of the process and leads to morphological abnormalities in the regenerated region. The severity in the delay in regeneration increases with the time period between anterior and posterior amputations. The number of worms with altered morphologies of the regenerated region increases with time after posterior amputation.

### Molecular characterization of posterior regeneration after both anterior and posterior amputations

To further understand defects in *P. dumerilii* posterior regeneration induced by anterior amputation, we took advantage of technics available nowadays and used various molecular markers. We performed whole-mount *in situ* hybridizations (WMISH) in C2 (anterior amputation two days before posterior amputation) and control (posterior amputation only) worms for nine genes whose expression during posterior regeneration was previously characterized (for more details on the nine studied genes, see Planques et al. 2019 and references therein). We chose C2 since worms in this condition were strongly delayed in their regeneration and, while displaying some abnormalities, had regenerated regions with morphologies that are not too severly altered, in order to be comparable with the controls. Representative images of the expression of the genes at five time points after posterior amputation (1, 2, 3, 5 and 7 dppa) are shown in Figs. 5 and 6.

**Figure 5.**
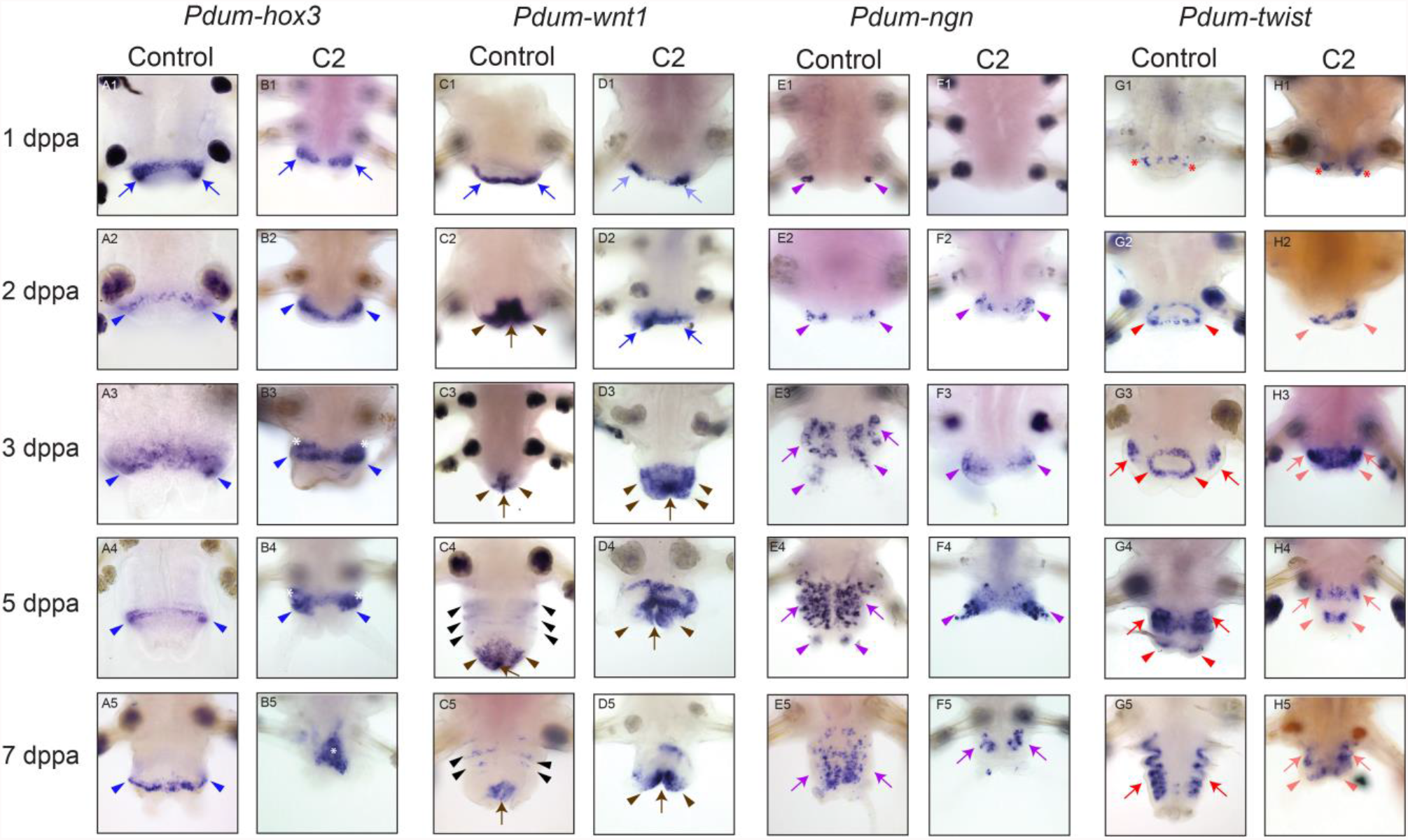
Effects of anterior amputation on the expression of genes involved in segment, organ or tissue patterning and differentiation during posterior regeneration. Whole-mount *in situ* hybridizations (WMISH) for the genes (n=4) whose name is indicated are shown for five time points (1, 2, 3, 5, and 7 days post-posterior amputation or dppa). Control worms are only posteriorly amputated. C2 worms are amputated anteriorly 2 days before posterior amputation. All panels are ventral views (anterior is up). Blue arrows = wound epithelium; blue arrowheads = posterior growth zone; white asterisks = extended or aberrant expression of *Pdum-hox3*; brown arrow = gut posterior-most part; brown arrowheads = pygidial epidermis; black arrowheads = segmental stripes; faint blue arrows = *Pdum-wnt1* restricted expression in the wound epithelium; purple arrowheads = putative sensory cell progenitors; purple arrows = developing ventral nerve cord; red asterisk = early expression in internal cells of *Pdum-twist*; red arrowheads = pygidial muscle cells progenitors; red arrows = segmental muscles; faint red arrowheads = misshapen expression domain of *Pdum-twist* in pygidial muscle cells progenitors; faint red arrows = misshapen expression domain of *Pdum-twist* in segmental muscles.

**Figure 6.**
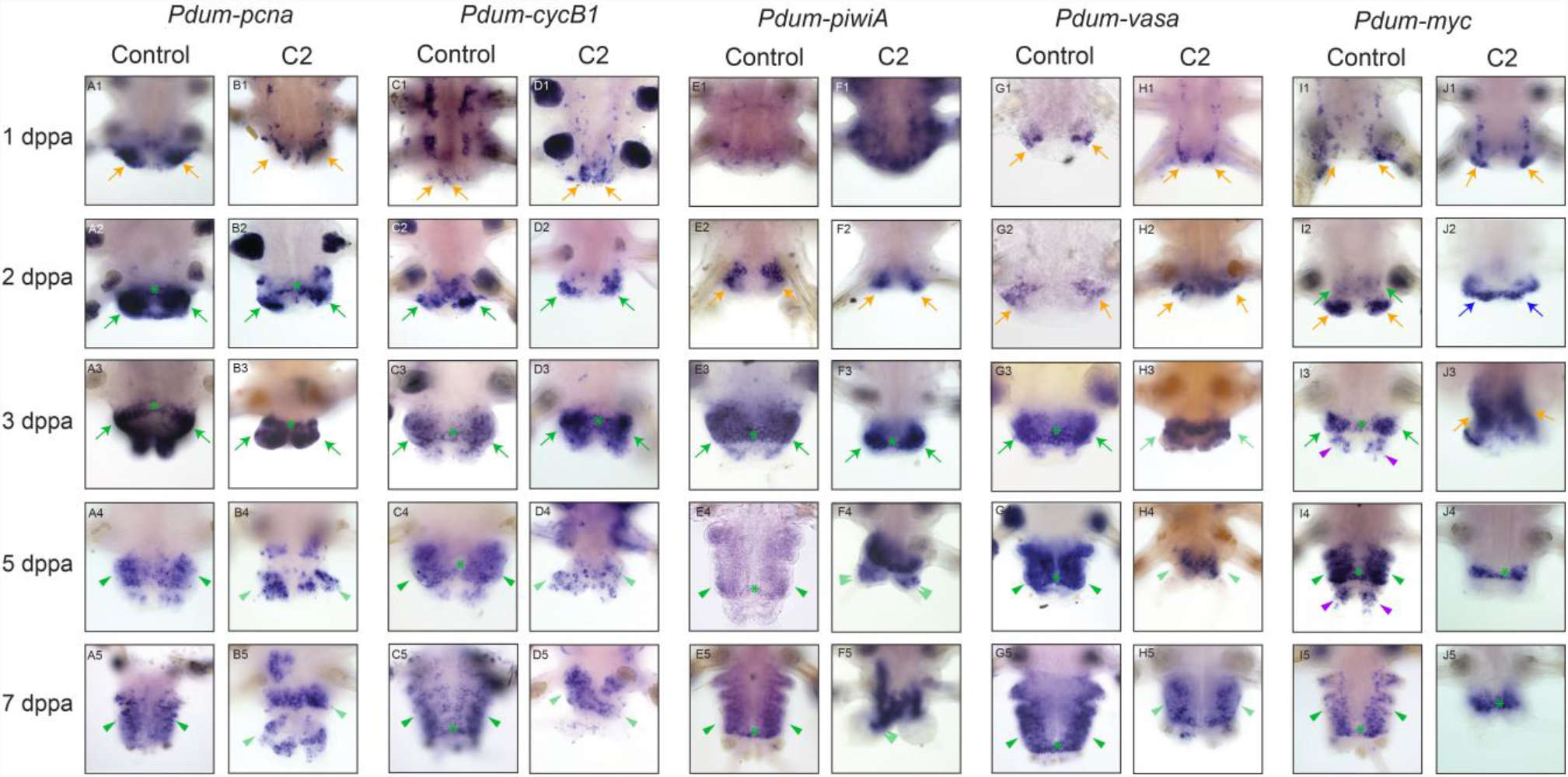
Effects of anterior amputation on the expression of cell proliferation and stem cells genes during posterior regeneration. Whole-mount *in situ* hybridizations (WMISH) for the genes (n=4) whose name is indicated are shown for five time points (1, 2, 3, 5, and 7 days post-posterior amputation or dppa). Control worms are only posteriorly amputated. C2 worms are amputated anteriorly 2 days before posterior amputation. All panels are ventral views (anterior is up). Orange arrows = proliferating internal cells below the wound epithelium; green arrow = large expression in the blastema; green arrowheads = large expression in the developing segments; green asterisks = growth zone stem cells; faint green arrowheads = reduced expression in the developing segments; double faint green arrowheads = aberrant expression in the pygidium; purple arrowheads = expression at the basis of the anal cirri; blue arrows = extended abnormal expression of *Pdum-myc*.

We first studied four genes whose expression allowed us to monitor the reformation of different structures and tissues during posterior regeneration. *Pdum-hox3* is a growth zone marker: this gene is first expressed in the wound epithelium at 1 dppa (Fig. 5A1, blue arrows) and, from 2 dppa onwards, in the growth zone stem cells (Fig. 5A2-A5, blue arrowheads). A similar expression was found in C2 worms at 1 and 2 dppa (Fig. 5B1, blue arrows; 5B2, blue arrowheads). At 3 and 5 dppa, a slightly extended and thicker expression domain of *Pdum-hox3* on the lateral sides of the regenerated region was observed in C2 worms (Fig. 5B3, B4; blue arrowheads and asterisks). A large asymmetrical expression, completely different to that found in controls, was observed at 7 dppa (Fig. 5B5; white asterisk). These data therefore suggest that, while probably initially regenerated in a more or less normal fashion, the growth zone is not properly maintained at later stages in C2 worms, which may explain why these worms do not produce new segments. *Pdum-wnt1* is a marker of the terminal structures of the worms (anus and pygidium). This gene is expressed at 1 dppa in the wound epithelium (Fig. 5C1; blue arrows), in the posterior-most part of the gut (Fig. 5C2-C5; brown arrows) and in the pygidial epidermis (Fig. 5C2-C5; brown arrowheads) from 2 dppa onwards, and in faint ectodermal stripes at 5 and 7 dppa (Fig. 5C4, C5; black arrowheads) which corresponds to segment primordia. In C2 worms, at 1 dppa, we found *Pdum-wnt1* expression to be restricted to the lateral part of the wound epithelium (Fig. 5D1; faint blue arrows). At 2 dppa, a strong expression in the wound epithelium was observed (Fig. 5D2; blue arrows), but in contrast to control animals no expression in the gut can be detected. At 3 dppa, the gene is expressed in a large posterior expression domain which appears extended when compared to controls and includes both epidermal and posterior gut cells (Fig. 5D3; brown arrowheads and arrows). At 5 and 7 dppa (Fig. 5D4, D5), *Pdum-wnt1* was expressed both in the posterior gut cells (brown arrow) and in the prospective pygidium, in an extended fashion (brown arrowheads). No epidermal segmental stripes of *Pdum-wnt1*-expressing cells were observed, confirming the impairment of segment production in C2 worms. *Pdum-wnt1* expression also suggests that the overall patterning of the regenerated region, notably the pygidium, is altered in C2 worms.

*Pdum-ngn* is a nervous system marker expressed in neural progenitors, first in a few lateral cells, which could be sensory cell progenitors, at 1 and 2 dppa (Fig. 5E1, E2; purple arrowheads), and, from 3 dppa onwards, in many neural precursors of the developing ventral nerve cord (Fig. 5E3-E5, purple arrows) and in sensory cells in the anal cirri (Fig. 5E3, E4, purple arrowheads). In C2 worms, *Pdum-ngn* expression was not detected at 1 dppa (Fig. 5F1) but was roughly normal at 2 dppa (Fig. 5F2, purple arrowheads). The number of *Pdum-ngn*-expressing cells was strongly reduced, as compared to controls, in later stages (Fig. 5F3-F5), strongly suggesting that nervous system formation is impaired in C2 worms in which some neurogenesis does however occur both in the anal cirri (Fig. 5F3, F4, purple arrowheads) and in the ventral nerve cord (Fig. 5F5, arrows). *Pdum-twist* is a muscle marker expressed in muscle progenitors and their differentiating progeny. It is weakly expressed in internal cells at 1 dppa (Fig. 5G1; red asterisks), then expressed in a ring of posterior cells at the origin of the pygidial muscles from 2 to 5 dppa (Fig. 5G2-G4; red arrowheads), and in segmental somatic muscles from 3 dppa onwards (Fig. 5G3-G5; red arrows). Expression of *Pdum-twist* appeared roughly normal in C2 worms at 1 dppa (Fig. 5H1, red asterisks). Expression in putative pygidial muscle precursors was observed at the next stages, in a pattern that however differs from that of control animals (absence of ring shape and/or extended expression, Fig. 5H2-H4, faint red arrowheads), suggesting that pygidial muscles are produced in an abnormal pattern in C2 worms. Expression of *Pdum-Twist* was also found in cells located more anteriorly in the regenerated region (Fig. 5H3–H5, faint red arrows), which may be somatic segmental muscles. However, unlike control worms, no segmental stripes of *Pdum-Twist* were observed. This therefore suggests that while some myogenesis occurs, muscle formation is nevertheless strongly altered in C2 worms.

We next studied the expression of two genes linked to cell proliferation, *Pdum-pcna* and *Pdum-cycB1* (Fig. 6A-D). At 1 dppa, *Pdum-pcna* is expressed in two large groups of cells close to the would epithelium (Fig. 6A1; orange arrows), while *Pdum-cycB1* is only expressed in a few cells in this same region (Fig. 6C1; orange arrows). From 2 dppa onwards, the two genes become largely expressed in the blastema (Fig. 6A2, A3, C2, C3, green arrows) and subsequently in the developing segments (Fig. 6A4, A5, C4, C5, green arrowheads). Expression in the growth zone stem cells is also visible from 2 dppa onwards (Fig. 6A2, A3, C3-C5, green asterisks). Until 3 dppa, no convincing alterations of the expression patterns of those two genes were found in C2 worms, although *Pdum-pcna* expression level, as roughly judged by labeling intensity, appeared reduced in these worms as compared to controls (Fig. 6B1-B3, D1-D3). From 5 dppa onwards, in contrast, the number of cells expressing the two genes was strongly reduced in C2, and their overall expression patterns are strikingly altered (Fig. 6B4, B5, D4, D5, faint green arrowheads). No expression in the growth zone stem cells can be detected in C2 worms. These data suggest that in C2 worms head amputation does not prevent cell proliferation during early stages of regeneration. In contrast, at later stages, proliferation is likely strongly reduced in C2 worms, in link with the failure to properly form new segments as shown in the previous sections.

We lastly studied the expression of three genes previously shown to be expressed in the stem cells of the growth zone, as well as in proliferating cells of the developing segments, *Pdum-piwiA, Pdum-vasa* and *Pdum-myc. Pdum-piwiA* becomes expressed at 2 dppa in two bilateral groups of internal cells (Fig. 6E2, orange arrows). Its expression extends to most blastemal cells at 3 dppa (Fig. 6E3, green arrows) and is detected in cells of the mesodermal part of the growth zone (Fig. 6E3, green asterisk). At 5 and 7 dppa, *Pdum-piwiA* continues to be expressed in the mesodermal growth zone (Fig. 6E4, E5, green asterisk), as well as in the mesodermal cells of the developing segments (Fig. 6E4, E5, green arrowheads). In contrast, no expression is found in the differentiating pygidium (Fig. 6E5). In C2 worms, no major abnormalities in the expression pattern of *Pdum-piwiA* were found until 3 dppa, although, as reported for *Pdum-pcna*, the expression level seemed to be reduced as compared to controls (Fig. 6F1-F3). At 5 and 7 dppa, *Pdum-piwiA* expression pattern in C2 worms was found to be strikingly different from controls, with no segmental expression and a broad intense expression including in the pygidium (Fig. 6F4, F5, double faint green arrowheads). No expression in mesodermal growth zone stem cells was detected. At 1 dppa, *Pdum-vasa* expression is restricted to two lateral patches of ectodermal cells (Fig 6G1, orange arrows) in control worms. From 2 dppa onwards, its expression is similar to that of *Pdum-piwiA* in control worms (Fig. 6G2-G5). In C2 animals, *Pdum-vasa* expression was found to be roughly normal at 1 and 2 dppa (Fig. 6H1, H2). At 3 dppa, no clear expression in the mesodermal growth zone stem cells is observed (Fig. 6H3), while a broad expression in the blastema, similar to that in controls is present (Fig. 6H3, green arrows). From 4 dppa onwards, the expression of the gene in C2 worms was strongly reduced as compared to controls, with no clear expression in the mesodermal growth zone stem cells (Fig. 6H4, H5, faint green arrowheads). In control worms, *Pdum-myc* is first expressed in a few scattered cells at 1 dppa (Fig. 6I1, orange arrows). It is strongly expressed at 2 dppa in two groups of posterior cells, at the position where anal cirri will form (Fig. 6I1, orange arrows) and weakly in more anterior blastemal cells (Fig. 6I2, green arrows). From 3 dppa onwards (Fig. 6I3-I5), *Pdum-myc* is expressed in the growth zone stem cells (Fig. 6I3–I5, green asterisk), in cells at basis of the anal cirri (Fig. 6I3, I4, purple arrowheads), and both in mesodermal and ectodermal cells of developing segments (Fig. 6I4, I5, green arrowheads). In C2 worms, at 1 dppa, *Pdum-myc* was found to be expressed in two small groups of lateral cells (Fig. 6J1, orange arrows). From 2 dppa onwards, we found that its expression pattern in C2 worms strikingly differed from that of controls (Fig. 6J2-J5). A ring of expressing cells was observed at 2 dppa (Fig. 6J2, blue arrows), a broad and diffuse expression at 3 dppa (Fig. 6J3, orange arrows), and at 5 and 7 dppa, an expression that seemed to be restricted to the growth zone and some adjacent cells (Fig. 6J4, J5, green asterisk). No segmental expression could be detected. The expressions of the three studied stem cell genes were therefore strikingly modified in C2 worms as compared to controls, albeit in different ways, and these alterations are suggestive of defaults in growth zone regeneration and subsequent segment formation.

In conclusion, the molecular characterization of the defects observed during *P. dumerilii* posterior regeneration following an anterior amputation highlights how the overall patterning of the regenerated regions is altered, especially in late regeneration stages. While some neurogenesis and myogenesis occur, the pygidium does not differentiate properly and the growth zone, while probably initially regenerated, does not function normally. This is in line with the fact that cell proliferation is highly reduced and stem cells markers are mis- and/or under-expressed. Altogether, this leads to the non-production of new segments and therefore to an impairment of the post-regenerative posterior growth process.

### Effects of anterior amputation on post-regenerative posterior growth

As mentioned before, previous studies reported that the removal of the brain impairs not only posterior regeneration but also post-regenerative posterior growth (*e*.*g*., Clark & Bonney 1960; Clark & Evans 1961; Clark & Scully 1964; Golding 1967c; Hofman 1976). To further evaluate the effects of brain removal on post-regenerative posterior growth in *P. dumerilii*, we performed a set of experiments combining posterior amputation followed by anterior amputation (Fig. 7A, red diamond and black diamond, respectively). Anterior amputations were performed after the regeneration of the growth zone which occurred between 2 and 3 dppa for 3-4-month-old worms (Planques et al. 2019). Three experimental conditions (for a total of 92 amputated individuals) plus one control experiment (n= 38; only posterior amputation) were designed for a total duration of ten days (Fig. 7A). For all experiments, day 0 was defined as the day when posterior amputation was performed. In condition C3, anterior amputation was performed 3 days post posterior amputation (3 dppa), there was thus a delay of three days between anterior and posterior amputations. Conditions C4 and C5 were designed similarly, with respectively four and five days of delay between posterior and anterior amputations. Worms were monitored every day for ten days and scored for the presence or absence of new segments produced by the regenerated growth zone. The overall results of these experiments are depicted in Fig. 7B. Detailed statistics can be found in Supp. Table 1. The morphology of the regenerated region was also monitored and abnormal individuals with morphologies difficult to assign to a precise stage were excluded from the main statistical analysis and their proportion was analyzed separately (Supp. Fig. 2).

**Figure 7.**
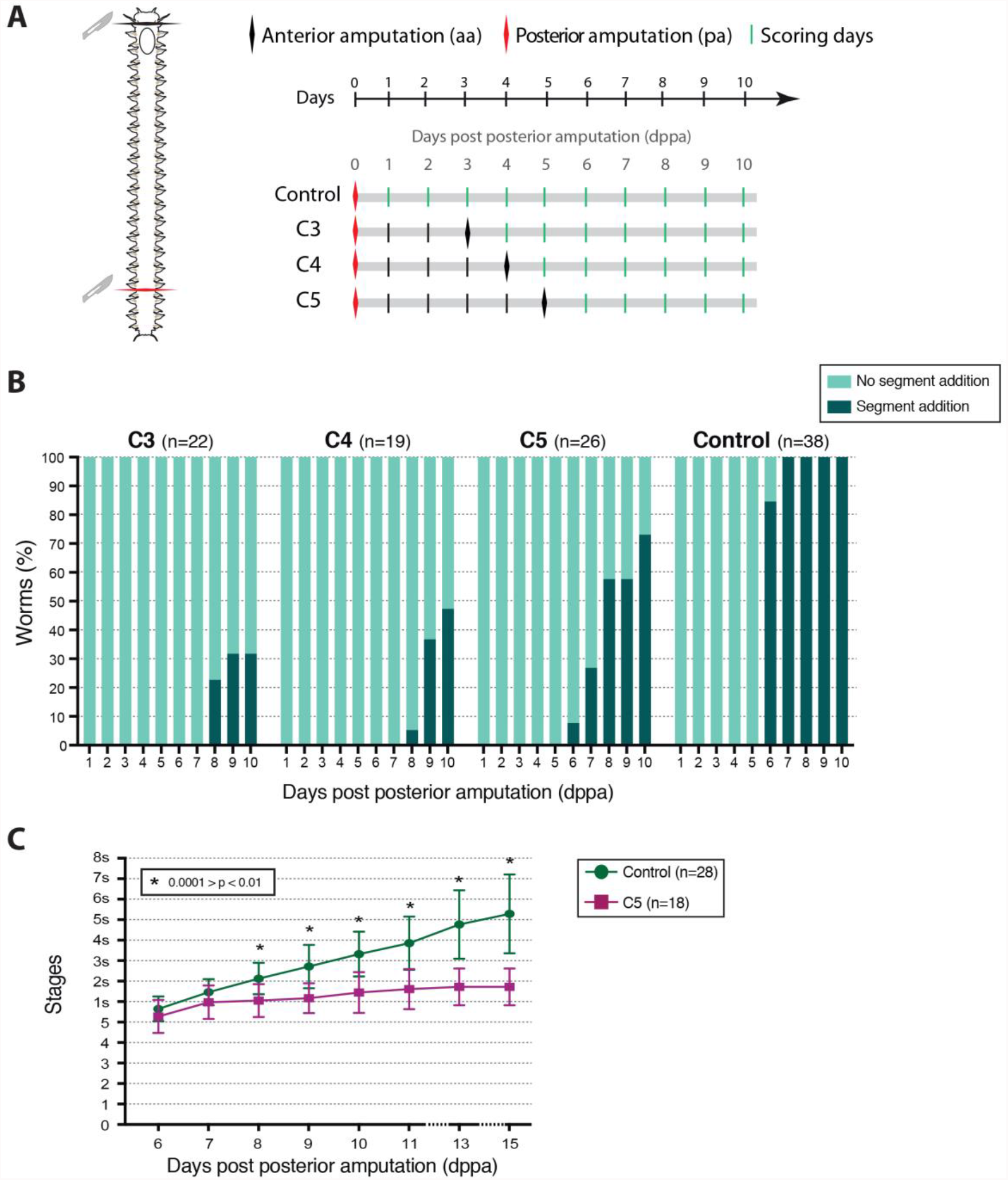
Influence of anterior amputation on *P. dumerilii* post-regenerative posterior growth. A) Schematic representation of the experimental design used to address the impact of anterior amputation (aa, black diamonds) on post-regenerative posterior growth (*i*.*e*., following posterior amputation or pa, red diamonds). A timeline representing the whole duration of the experiment (10 days) and the timepoints for anterior amputation and posterior amputation for each condition, as well as the scoring days (green bars, from 1 to 10 days post posterior amputation, dppa), are provided. Control worms were only amputated posteriorly; for C3, anterior amputation was done 3 days after posterior amputation; similarly, for C4 and C5, anterior amputations were done 4 and 5 days after posterior amputation, respectively. B) Graphic representation of the percentage of worms that have undergone post-regenerative posterior growth (addition of at least one new segment) every day for 10 days after posterior amputation, for each condition (the number of scored individuals is indicated next to the condition name). C) Graphic representation of the regeneration stages reached by the worms or the number of newly-added segments (1s = 1 segment, 2s = 2 segments, …) every day from 6 dppa to 15 dppa in control (only posterior amputation) and C5 (anterior amputation five days after posterior amputation) worms (see inset for the color code and the number of worms used per condition). Mean + SD are shown. Asterisks indicate significant differences as defined by 2-way ANOVA on repeated measures with Dunnett correction. * 0.0001 > p < 0.01 (for specific p-values, see Supp. Table 1).

As expected, control worms show no sign of segment addition between 1 to 5 dppa (Fig. 7B and Supp. Table 1). At 6 dppa, the great majority of control worms (84,6%) present new segment(s) produced by the regenerated growth zone (Fig. 7B). From 7 dppa onwards, all control worms have added at least one new segment (Fig. 7B). C3, C4 and C5 worms show no sign of segment addition between 1 to 5 dppa (Fig. 7B). In striking contrast to control worms, C3, C4 and C5 worms showed a much-delayed posterior growth in subsequent stages. For the C3 condition, no new segment was produced at 6 and 7 dppa (Fig. 7B). A small proportion of C3 worms showed at least one segment from 8 dppa onwards (from 22.7% at 8 dppa to 31.8% at 10 dppa). Similarly, C4 worms present at least one new segment from 8 dppa onwards (Fig. 7B). While the proportion of worms showing post-regenerative posterior growth at 8 dppa is extremely reduced (5.3%), this percentage increases substantially at 9 and 10 dppa (36.8 % and 47.4 %, respectively). In contrast to C3 and C4, C5 worms present at least one segment from 6 dppa onwards (Fig. 7B). While a small percentage (7.7%) of worms have produced at least one segment at 6 dppa and few more (26.9%) at 7dppa, the majority of them present post-regenerative posterior growth at 8-10 dppa (57.7% at 8 and 9 dppa, and 73.1% at 10 dppa). For all conditions, significant differences, compared to the control, started to be observed one day after the anterior amputation was performed, *i*.*e*., 4 dppa in C3, 5dppa in C4 and 6 dppa in C5 (Supp. Table 1). The proportion of worms that were able to produce new segments and the timing of the beginning of post-regenerative posterior growth is thus highly correlated with the moment when the anterior amputation was performed. We also observed some worms with abnormal morphologies from 6 dppa onwards in C3 and C4 conditions (but not in C5 condition), corresponding to 18.75% and 10% of the worms in C3 and C4, respectively (Supp. Fig. 2A, B). At 7 dppa, worms with morphological abnormalities were observed in the three conditions, in a proportion of ∼18.75% in C3, ∼16.6% in C4, and ∼6.6% in C5 (Supp. Fig. 2A, B). At the end of the experiment (10 dppa), the proportion of worms showing abnormal morphologies increased mainly in C3 and C4, with ∼31% and ∼36% (respectively), whereas C5 showed only a ∼13% of them (Supp. Fig. 2A, B). No abnormalities were observed in the control worms.

In order to better characterize the impact of anterior amputation on post-regenerative posterior growth, we performed an additional experiment in which we assessed the number of new segments that have been produced (Fig. 7C). We first performed posterior amputations on 3-4-month-old worms and let these worms regenerate until 5 dppa. At this time point, we selected worms that had reached the regeneration stage 5 and then performed anterior amputation (C5 condition worms n=18) or no additional amputation (control worms n=28). We then scored both C5 and control worms for the number of newly-added segments until 15 dppa. At 6 and 7 dppa, no significant differences between C5 and control worms were observed, and the worms had produced an average of 0.5 and 1 segment, respectively (Fig. 7C). From 8 dppa onwards, significant differences between C5 and control worms were observed: C5 worms only added one more segment during the seven following days (until 15 dppa), while control animals produced an average of five segments during the same time period (Fig. 7C). Unexpectedly, no abnormal morphologies were found until 11 dppa, when a proportion of ∼11% of the worms showing morphological abnormalities was observed, a proportion that reached ∼33% at the end of the experiment at 15 dppa (Supp. Fig. 2C, D). No abnormalities were observed in the control worms.

Altogether, these results show that after anterior amputation post-regenerative posterior growth is drastically affected and that a very limited number of new segments is produced. To further characterize these defects, we performed WMISH on C5 and control worms at 10 dppa for *Pdum-pcna* to monitor cell proliferation, *Pdum-hox3* to visualize the stem cells of the regenerated growth zone, and *Pdum-en* (*Pdum-engrailed*) to follow the production of new segments and therefore the activity of the growth zone (Fig. 8). As expected, control and C5 worms harbor a quite different morphology, as very few post-regenerative segments have been produced in C5 worms compared to the control ones. In control worms, *Pdum-pcna* is broadly expressed in the developing segments (Fig. 8A, green arrowheads). In C5 worms a similar expression is observed, yet in a smaller region given the reduced number of newly produced segments (Fig. 8A’, green arrowheads). The ring-shaped expression of *Pdum-hox3* in the growth zone located at the anterior margin of the pygidium, found in the control worms (Fig. 8B, blue arrowheads), is also present in the C5 worms (Fig. 8B’, blue arrowheads), suggesting the presence of the growth zone in the C5 worms. Finally, while several segmental stripes of *Pdum-en* expression delineating the newly produced segments are found in the control worms (Fig. 8C, brown arrowheads), only one or two such stripes are present in the C5 worms (Fig. 8C’, faint brow arrowheads), suggesting a defect in the production of segments by the growth zone and therefore alterations of the function of the growth zone stem cells in C5 worms.

**Figure 8.**
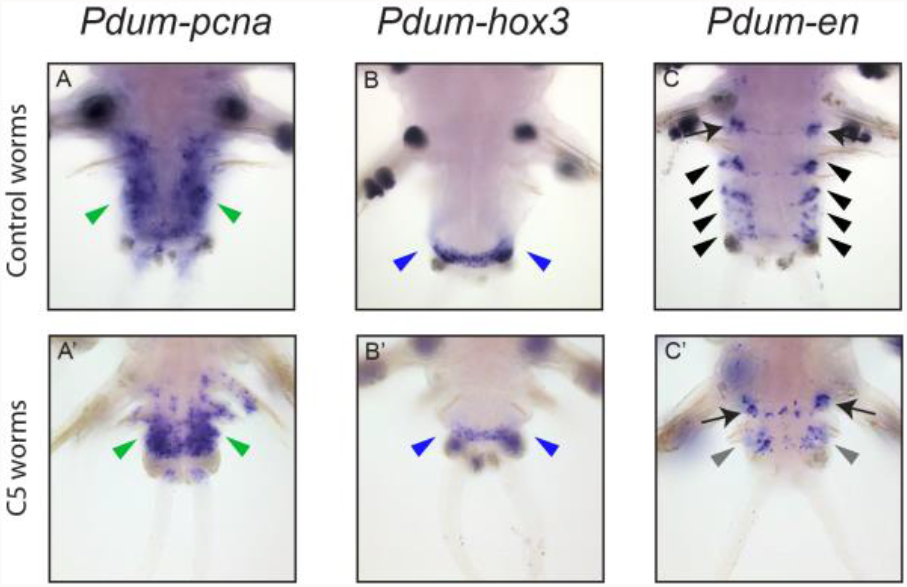
Effects of anterior amputation on the expression of cell proliferation, growth zone and segment marker genes during post-regenerative posterior elongation. Whole-mount *in situ* hybridizations (WMISH) for the genes (n=3) whose name is indicated are shown at ten days post-posterior amputation (10 dppa) for C5 (anterior amputation five days after posterior amputation) and control (no anterior amputation) worms. All panels are ventral views (anterior is up). Green arrow = large expression in the regenerating structure; blue arrowheads = posterior growth zone; black arrowheads = segmental stripes; faint black arrowheads = abnormal segmental stripes in a reduced number; black arrows = expression at the border between the last non-amputated segment and the regenerated region.

In conclusion, we have shown that after anterior amputation post-regenerative posterior growth is deeply affected, and the production of post-regenerative segments is extremely limited. These drastic defaults are not due to the absence of the growth zone as shown by the presence of *Pdum-hox3*-expressing cells at 10 dppa (*i*.*e*., five days after anterior amputation), but are likely due to an altered functioning of those cells.

### Effects of exogenous methylfarnesoate (MF) on posterior regeneration after both anterior and posterior amputations

Methylfarnesoate (MF) has been reported to likely be the putative brain hormone controlling sexual maturation and suggested to also be involved in the control of posterior regeneration and growth in *P. dumerilii* (Schenk et al. 2016). We have shown (see previous sections) that, while not affecting early steps of posterior regeneration, the amputation of the head prevents regeneration from being completed and leads to morphological abnormalities in the regenerated region. To assess whether the defects in posterior regeneration after head amputation are due to the absence of MF, we tested if these defects can be rescued by the exogenous addition of MF hormone, as it was the case when supraoesophageal ganglia from intact worms were implanted in decerebrated host (*e*.*g*., Scully 1964; Golding 1967b). As in the previous experiments, we used 3-4-month-old worms and compared three sets of worms. Worms of the “control 1” condition (n=29) were amputated only posteriorly at day 0 and display a normal posterior regeneration process; worms of the “control 2” condition (n=27) were posteriorly and anteriorly amputated at day 0, and present delay and defects in posterior regeneration (these worms correspond to the C0 condition in Figure 3); MF-treated worms (n=51) were similarly posteriorly and anteriorly amputated at day 0 but were subsequently incubated in sea water containing 100 nM of MF hormone, renewed every 24 hours (Fig. 9A). The three sets of worms were scored for the regeneration stage that was reached every day for five days (until 5 dppa). The overall results of these experiments are depicted in Fig. 9B. Morphology of the regenerated region was also monitored (Supp. Fig. 3) and abnormal individuals with morphologies difficult to assign to a precise stage were excluded from the statistical analysis.

**Figure 9.**
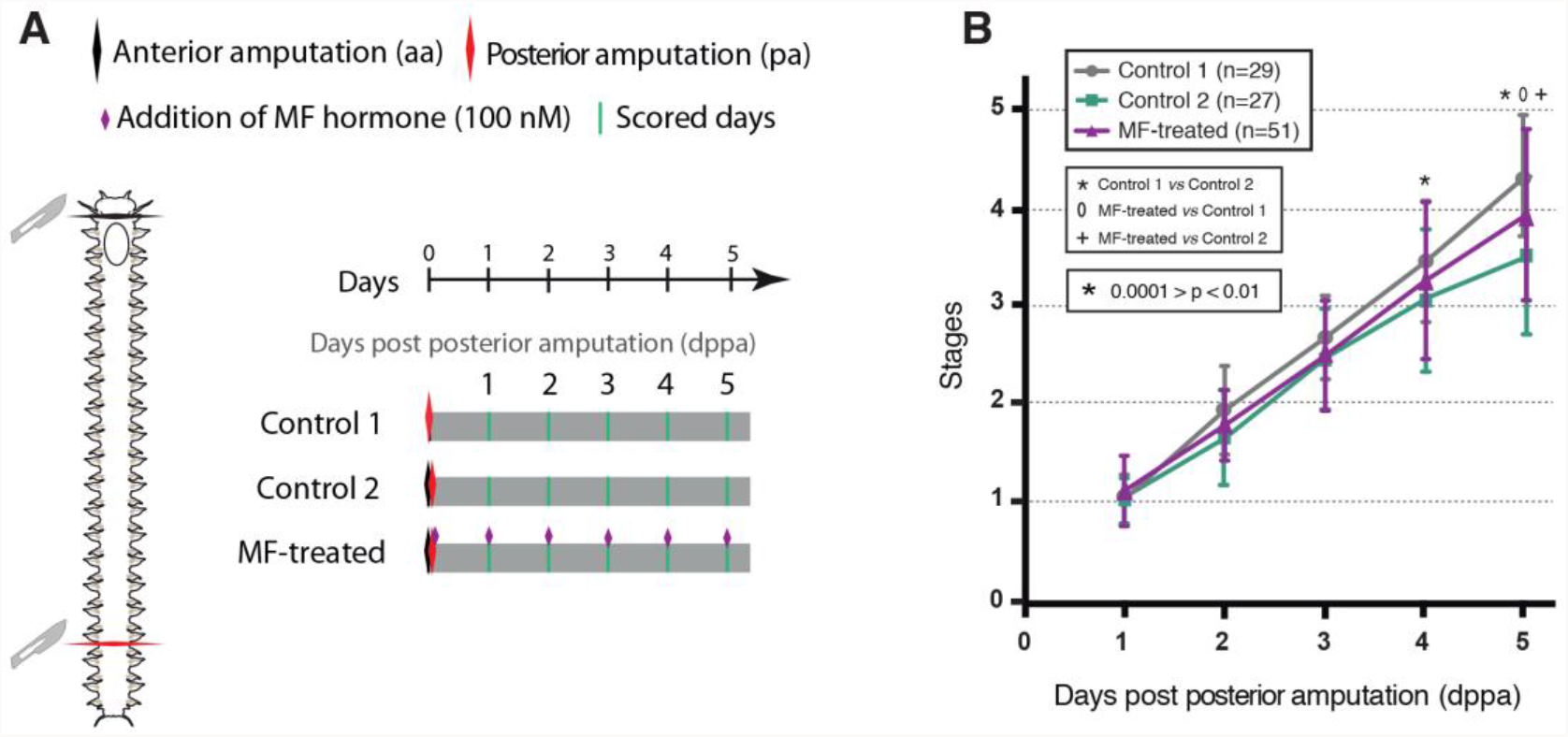
Assessment of the effect of methylfarnesoate (MF) on *P. dumerilii* posterior regeneration. A) Schematic representation of the experimental design used to address the impact of exogenous MF treatment on posterior regeneration. A timeline representing the whole duration of the experiment (5 days) and the timepoints for anterior amputation (aa) and posterior amputation (pa) for three conditions, as well as the scoring days (green bars, from 1 to 10 days post posterior amputation, dppa), are provided. Control 1: posterior amputation, no anterior amputation, no MF treatment; Control 2: posterior amputation, anterior amputation, no MF treatment; MF-treated: posterior amputation, anterior amputation, MF treatment (100nM in sea water, renewed every day). B) Graphic representation of the regeneration stages reached by the worms every day for 5 days after posterior amputation, for each condition (see inset for the color code and the number of worms used per condition). Mean + SD are shown. Significant differences (0.0001 > p < 0.01) are indicated (see inset). Statistics: 2-way ANOVA on repeated measures with Tukey correction.

As expected, from 4 dppa onwards, control 2 worms were significantly delayed as compared to the control 1 condition (Fig. 9B and Supp. Table 1). Interestingly, at 4 dppa, MF-treated worms were not significantly different to neither control 1 nor control 2 worms. At 5 dppa, however, MF-treated worms were significantly delayed compared to the control 1 worms but also regenerated significantly faster than the control 2 worms (Fig. 9B and Supp. Table 1). These results therefore indicate that the addition of exogenous MF positively affect posterior regeneration in headless worms. However, as MF-treated worms did not regenerate as the control 1 worms, it also indicates that MF is not completely reverting the defect produced by head amputation. We observed a similar proportion of worms with abnormal morphologies both in control 2 (13%) and MF-treated worms (15%) (Supp. Fig. 3 and Supp. Table 1), further indicating that the treatment with exogenous MF cannot fully compensate for the absence of the head.

## Discussion

### Brain hormonal activity is required for both posterior regeneration and post-regenerative posterior growth

Limits to the energy that all living organisms can expend to maintain their structure and to carry out their main regulatory processes entail the decision on how to best invest the energetic resources available. This is especially crucial for semelparous species that only reproduce once in their life and die afterwards, and therefore need to reduce the investment in their somatic growth or other dispensable activities when they become sexually mature (*e*.*g*., Kozlowski & Wiegert 1986; Stearns 1992; Bonnet 2011). Within the marine realm, nereids may be one of the best examples to observe the costs and benefits of this strategy, given that they reproduce through epitoky. Epitoky involves drastic morphological, physiological, and behavioural changes required for reproduction and, as a counterpart, induces the loss of their capacity to grow and regenerate missing tissues after an injury (*e*.*g*., Hauenschild & Fischer 1962; Golding 1967a, 1972; Landau et al. 2010).

It is well established that this switch between sexual reproduction *versus* growth/regeneration is mediated by an endocrine control from the brain (specifically by the supraoesophageal ganglion). Indeed, several studies showed that the removal of this ganglion promotes sexual maturation and inhibits posterior growth and regeneration (*e*.*g*., Durchon 1952; Casanova 1955; Clark & Bonney 1960; Clark & Evans 1961; Clark & Ruston 1963; Durchon & Marcel 1962). In addition, the implantation of a supraoesophageal ganglion (taken from a young worm) in the coelom of headless old animals that were already starting their sexual metamorphosis, was shown to block epitoky and the maturation of gametes. On top of that, this experimental procedure also allowed worms that initially were no longer able to regenerate to recover the capability to regenerate upon amputation (Durchon 1952; Hauenschild 1956a; Hauenschild & Fischer 1962). In addition, if the ganglion from the donor was subsequently removed from the host, the metamorphosis to sexual mature individuals resumed and regenerative abilities were lost (Hauenschild & Fischer 1962). Furthermore, the onset of normal metamorphosis can be delayed in intact worms (non-decerebrated) by the implantation of immature ganglia (Clark & Ruston 1963). From all these experiments, it has been hypothesized that the supraoesophageal ganglion produces a hormone(s) that represses sexual maturation and promotes growth and regeneration, and that hormone production is at its highest level in young worms and progressively declines when the worms become older (Clark & Scully 1964; Scully 1984).

Although there is no doubt, from the aforementioned studies mostly conducted in the fifties and sixties, about the important role that the brain and its hormonal production play in the reproductive and regenerative processes of nereid annelids, it is important to note that all these studies usually showed some limitations. Firstly, animals were typically collected from the wild, and therefore, a proper estimation of their age was difficult to assess. The number of segments, and therefore the size of the worms, increases with the age, but there is no strict correlation between the number of segments, the overall size and the age of the worms (as the density of worms has also an impact on their size and segment number). Age, number of segments, and size can all influence the overall regeneration process, notably its speed as shown by our recent study performed on worms raised in laboratory conditions, which demonstrated that regeneration was significantly faster in small worms with 10-20 segments (1 month old) than in older worms displaying 30-40 or 70-80 segments (Planques et al. 2019). It is also likely that the conditions in which the worms are raised (for example nutrition or temperature) may influence their regeneration, as they influence development and growth (Kuehn et al 2019; Fischer et al 2010), further underlining for the importance of using standardized laboratory conditions. Secondly, another important problem of most reported studies was the lack of distinction between posterior regeneration and post-regenerative posterior growth (*e*.*g*., Golding 1967a). The occurrence of regeneration was usually established based on the presence of newly added segments (Clark & Bonney 1960; Clark & Evans 1961; Clark & Ruston 1963), which may lead to biases in the conclusions drawn. We previously argued that a clear distinction should be made between posterior regeneration *per se*, which involves the restoration of the pygidium and the posterior growth zone, and post-regenerative posterior growth, which comprises the generation of new segments from the regenerated posterior growth zone (Gazave et al. 2013; Planques et al. 2019). The regeneration phase consists of five well-defined and reproducible stages that follow a constant timeline, as long as worms homogenous in size and age are studied (Planques et al. 2019). Post-regenerative posterior growth is, in contrast, much more variable in term of number of segments that have been added at a given time point in different individuals (Planques et al. 2019). Lastly, the aforementioned studies showing the importance of the brain in controlling regeneration were made in the “pre-molecular” era and the lack of molecular data clearly limited the understanding of the regeneration defects that were observed after heads amputation or brain removal.

The recent use of molecular and cellular markers has made an in-depth characterization of both normal posterior regeneration and post-regenerative posterior growth possible (Gazave et al. 2013; Planques et al. 2019) and provides invaluable tools to further characterize the role of the brain during both processes. We therefore decided to re-assess and further analyze the role of the brain hormonal activity on posterior regeneration and post-regenerative posterior growth in *P. dumerilii* (i) using homogenous worms in terms of size and age and raised in stereotypical laboratory conditions and (ii) combining morphological and molecular data to increase our understanding of this role. We performed anterior amputations that remove the brain, at different time points before or after posterior amputation, to assess their effects on posterior regeneration and post-regenerative posterior growth, respectively.

#### Effects on posterior regeneration

To investigate the effect of anterior amputation on posterior regeneration, we first performed anterior amputations either at the same time than posterior amputation (condition C0; Fig. 3A) or at six different time points (from one day to seven days) before posterior amputation (conditions C1 to C7; Fig. 3A). We found that regeneration was impaired in all conditions since, in contrast to control worms, no worms whose head was amputated were able to complete regeneration (*i*.*e*., to reach stage 5) at 5 and even 7 days post posterior amputation (dppa) (Fig. 3B). Consequently, none of the decapitated worms were able to add new segments. Morphological abnormalities, never observed in controls, were also found in worms of all conditions (Fig. 4). These results confirm the absolute requirement of the head (and thus likely the brain) for proper posterior regeneration. A second important observation is that the severity of the delay in posterior regeneration increases with the time period between anterior and posterior amputations. The least substantial delays were found for C0 and C1 conditions, where some worms were able to reach stage 4, while the strongest delays were found in C5, C6 and C7 conditions, where none of the worms reach this stage and most or all of them were even unable to reach stage 3 (Fig. 3B). These data are therefore consistent with the hypothesis that anterior amputation impairs posterior regeneration by causing a decrease in the concentration of the brain hormone in the blood and/or tissues of the worms, due to the absence of the brain where this hormone is produced. Significant differences in the regeneration stages reached by the worms were observed between C0 and C1 conditions, on the one hand, and all other conditions (C2-C7) on the other hand (Supp. Table 1), suggesting a rapid decrease in the hormone concentration in the absence of its production by the brain. A third compelling observation is that in all conditions, headless worms were able to reach stage 1 and stage 2. Stage 1 corresponds to the completion of wound healing, and, at stage 2, a small blastema is formed and the anus is regenerated (Planques et al. 2019). These early steps of the regeneration process therefore likely occur irrespective of the presence or absence of the brain hormone. Further progression through the following steps of regeneration, in contrast, does not happen in the absence of the head, therefore suggesting that these later steps require the activity of the brain hormone. Treatments with the cell proliferation inhibitor hydroxyurea similarly showed that stage 1 and 2 can be reached in the absence of cell divisions but that the following steps were cell proliferation-dependent (Planques et al. 2019), raising the possibility that head amputation may affect regeneration though an effect on cell proliferation.

To further characterize the defects in the posterior regeneration process caused by anterior amputations, we analyzed the expression of several genes that are well-known markers of specific tissues/structures during posterior regeneration in C2 worms (Fig. 5). Several conclusions can be drawn from this molecular analysis. Firstly, the posterior growth zone is initially regenerated in C2 worms as seen by the proper expression of *Pdum-hox3* (a growth zone marker) at early time points of posterior regeneration (1 to 2 dppa, Fig. 5 A1-2 *vs* B1-2). Interestingly, the regenerated growth zone is not able to produce any segments as seen by the absence of visible segments but also the absence of stripes of cells expressing *Pdum-wnt1* (a marker of segmentation) at 5 and 7 dppa (Fig. 5, C4-5 *vs* D4-5). From 3 to 5 dppa, the growth zone starts to show some defaults, being enlarged in C2 worms as shown by a large row of cells expressing *Pdum-hox3* (Fig. 5 A3-4 *vs* B3-4). At 7 dppa, the expression of *Pdum-hox3* in C2 worms becomes very different from that in controls (Fig. 5 A5 *vs* B5), suggesting an improper maintenance of the regenerated growth zone in the absence of the head. The expression of *Pdum-Wnt1* highlights the fact that other structures of the regenerating posterior part are also severely affected in C2 worms. Indeed, the wound epithelium appears to be incompletely formed (Fig. 5 C1 *vs* D1) and the pygidium morphology is extremely affected from 3 dppa onwards (Fig. 5 C3-5 *vs* D3-5). Several other aspects of posterior regeneration are also altered in C2 worms as shown by the strong reduction or misshape of the expression domain of *Pdum-twist* and *Pdum-neurogenin* (Figure 5 E, F, G, and H), which indicates that the formation of muscles and the nervous system, respectively, are impaired in the absence of the head. Taken together, these observations strongly suggest that the brain hormonal activity is required for many aspects of posterior regeneration including proper regeneration of the growth zone (and its subsequent ability to produce segments), wound epithelium and pygidium formation, as well as muscle and nervous system regeneration.

Secondly, the normal expression of the cell cycle genes *Pdum-cycB1* and *Pdum-pcna* at early stages of posterior regeneration in C2 worms (until 3 dppa; Fig. 6 A1 to D2) suggests that head amputation does not block cell proliferation during these early stages. It also suggests that the defects caused by head amputation are not primarily due to the blocking of cell proliferation but rather to a patterning default. Accordingly, the expression patterns of *Pdum-hox3, Pdum-twist*, and *Pdum-neurogenin* in C2 worms clearly differ from those of these genes in worms that have been treated with the cell proliferation inhibitor hydroxyurea after posterior amputation (Planques et al. 2019), indicating that head amputation and the inhibition of cell proliferation have distinct effects on posterior regeneration. At 5 and 7 dppa, the expression of *Pdum-cycB1* and *Pdum-pcna* are strikingly reduced as compared to control animals, suggesting a decrease in cell proliferation at these later stages in C2 worms, which might be due to the failure to produce segments (Fig. 6 A4 to D5). Abnormal expression patterns of three stem genes previously shown to be expressed in proliferative cells, *Pdum-piwiA, Pdum-vasa*, and *Pdum-myc* were also observed in C2 worms at these late time points, confirming major defects in the pattern of proliferative cells causes by head amputation.

#### Effects on post-regenerative posterior growth

To investigate the effect of anterior amputation on post-regenerative posterior growth, we first performed anterior amputations 3, 4, or 5 days after posterior amputation (conditions C1, C3, and C5; Fig. 7A). These three conditions were chosen because we previously showed that the growth zone was regenerated and functional at three days after posterior amputation (Planques et al. 2019). We first checked the occurrence of post-regenerative posterior growth in the worms of these three conditions, as compared to control animals that were only posteriorly amputated. While most control worms added at least one new segment at 6 dppa and all of them from 7 dppa onwards, we observed a striking reduction of the number of worms undergoing post-regenerative posterior growth from 6 dppa to 10 dppa in all three experimental conditions (Fig. 7B). This reduction was dependent on the time point when the anterior amputation has been performed: at 10 dppa for example, about 30% of the worms showed post-regenerative posterior growth in C3, about 45% in C4, and about 75% in C5. Post-regenerative posterior growth is therefore clearly affected by anterior amputation. Two additional observations can be done: (i) the defects are more severe if the anterior amputation is done earlier (C3>C4>C5); (ii) in all conditions, some of the decapitated worms were able to produce at least one new segment. These observations can be explained by the fact that the regenerated growth zone has already produced one segment primordium at 3 dppa and two at 5 dppa. Indeed, our previous data showed the presence at 3 dppa of one stripe of cells expressing the early segmental marker *Pdum-engrailed* and that a second stripe of *Pdum-engrailed*-expressing cells are observed at 4 and 5 dppa (Planques et al. 2019). Some variability among the worms does however exist, as not all individuals show this *Pdum-engrailed*-expressing cell stripe at 3 dppa. In the C3 condition, it therefore means that anterior amputation has been done in worms that have not all produced a segment primordium or whose segment that has been produced is still in a very early step of its formation. This would explain why most of the C3 worms do not show addition of visible segments at later time points. In the two other conditions, especially C5, head amputation is performed in worms that have already one or two segment primordia produced at earlier time points, which would explain why more worms display post-regenerative posterior growth at later time points such as 10 dppa.

We further characterize the C5 condition and observed that these worms, which we scored until 15 dppa, form at most two segments while control animals added an average of five segments for the same period of time (Fig. 7C). This strongly suggest that the C5 worms do not produce any segments after head amputation and therefore that the head, and thus the brain hormone is required for post-regenerative posterior growth. Accordingly, only one or two stripes of *Pdum-engrailed* were observed in C5 worms at 10 dppa (Fig. 8). Our data also suggest that segments whose primordium has been produced before head amputation may still differentiate in the absence of the head, suggesting that the brain hormone would be required to produce the segment and/or early steps of its development, but not for its full differentiation. Alternatively, the potential remaining of low quantities of brain hormone in the body during the few days that follow head amputation might be sufficient to allow the differentiation of segments produced by the regenerated growth zone before head amputation, but not to sustain the formation of new segments after head amputation. In addition, the normal expression of *Pdum-hox3* in C5 worms at 10 dppa suggests that the defect in segment formation is not due to the disappearance or degeneration of the growth zone but rather in some alterations of its functioning.

### Involvement of methylfarnesoate (MF) in posterior regeneration

A wealth of old studies has led to the hypothesis that the brain hormonal activity, which would consist of a single hormone being continuously secreted by the brain until sexual maturation, would exert both a positive control on posterior growth and regeneration and a negative control of sexual maturation (*e*.*g*., Durchon 1952; Clark & Ruston 1963; Durchon & Marcel 1962; Scully 1964; Hauenschild 1956a; Golding 1983; Golding 1967a; Hofman 1976). Indeed, the removal of the supraesophageal ganglion in juvenile *P. dumerilli* worms has been shown to both impair their capability to grow and regenerate and induce their sexual maturation (Hauenschild & Fischer 1962; Hauenschild 1956b). Both the loss of posterior growth/regeneration abilities and the induced sexual maturation could be reverted by implanting the supraesophageal ganglion of young worms in the decerebrated worms (Hauenschild & Fischer 1962). While the molecular nature of the brain hormone has remained elusive and controversial for many years, a recent study has identified methylfarnesoate (MF) as a most likely candidate for being the *P. dumerilii* brain hormone (Schenk et al. 2016). MF was convincingly shown to affect vitellogenesis, a key process for oocyte growth and maturation, as shown by its negative action on the production of the yolk precursor Vitellogenin by coelomic cells both *in vitro* and *in vivo* (Schenk et al. 2016). These results therefore strongly suggest that MF acts as a repressor of sexual maturation as expected for the brain hormone. In addition, it was also reported that body segments supplemented with exogenous MF showed enhanced expression of *Pdum-hox3* as compared to control segments not treated with MF, leading to the suggestion that MF may also act on posterior regeneration (Schenk et al. 2016). Additional experiments were however required for further support the involvement of MF in posterior regeneration in *P. dumerilii* and test whether MF mediates the role of the brain in this process.

To tackle these questions, we tested whether the exogenous addition of MF hormone might rescue the impairment of posterior regeneration caused by the absence of the head and consequently of the brain hormone. Worms that were both anteriorly and posteriorly amputated at the same time were incubated for 5 days in sea water containing MF (regeneration in the absence of the head, MF treatment) and the regeneration of these worms was compared to that of two types on control animals: worms that were only amputated posteriorly and with no addition of MF in sea water (control 1, regeneration in the presence of the head, no MF treatment), and those that were amputated both anteriorly and posteriorly and with no addition of MF in sea water (control 2, regeneration in the absence of the head, no MF treatment) (Fig. 9A). At 5 dppa, MF-treated worms regenerated significantly faster than control 2 animals but significantly slower than control 1 worms (Fig. 9B), indicating that the exogenous addition of MF was able to rescue the defects caused by the absence of the head, albeit only partially. These results provide a clear support to the hypothesis that MF can mediate the positive control exerted by the brain on posterior regeneration, reinforcing the previous claim that MF is the brain hormone and that it can act on both regeneration and sexual maturation (Schenk et al. 2016).

The rescue of the regeneration defects caused by head amputation that we obtained with MF treatments was however only partial. This is different from what has been obtained in previous studies involving the implantation of supraoesophageal glands in decerebrated worms, which were described as leading to a seemingly complete capacity to regenerate (*e*.*g*., Golding 1967b). This discrepancy could be due to the experimental design of the experiment, as MF is provided from the sea water in which the worms are incubated and not directly in the internal medium of the animals. We do not know what the stability of MF in sea water is, nor what the efficiency of the uptake of the compound by the worm cells is. It is therefore possible that the concentration of active MF in the internal medium of the worms in our experimental conditions is inferior to that in non-decapitated worms or decerebrated worms in which a supraoesophageal ganglion has been grafted. One possible explanation would therefore be that the amount of MF present in the body might be not high enough to completely revert the defects in posterior regeneration caused by the lack of the brain. We used the same concentration of MF in sea water than that used by Schenk et al. (2016) and which was shown to have a strong negative effect on Vitellogenin production in whole worms. It is however conceivable that different quantities of MF may be required for the different functions in sexual maturation and regeneration that this hormone fulfills. We may also not exclude that the brain hormonal activity may involve (an)other hormone(s) than MF and therefore that supplementing MF alone is insufficient to compensate for the absence of the brain and rescue the defects in the posterior regeneration process due to this absence.

## Conclusions

We re-addressed the role of the brain hormonal activity on posterior regeneration and post-regenerative posterior growth in the annelid *P. dumerilii*. We show that the removal of the head prevents posterior regeneration from being successful and impairs multiple aspects of this regenerative process. We also show that post-regenerative posterior growth is severely affected in the absence of the brain. Indeed, while the posterior growth zone is apparently regenerated, its functioning is affected in decapitated worms. Our results therefore strongly suggest a key requirement of brain hormonal activity for regeneration and growth in *P. dumerilii*. We also show that exogenous methylfarnesoate (MF), which was proposed to be the *P. dumerilii* brain hormone and shown to act on worm reproduction, partially rescues posterior regeneration defects in decapitated worms, supporting the hypothesis that MF mediates the positive control of the brain on posterior regeneration. Our study therefore extends our knowledge about the control of posterior regeneration by the brain in *P. dumerilii*, paving the way for future mechanistic analysis of hormonal control of regeneration in this species.

## Acknowledgements

We thank all Vervoort lab members for helpful discussions and feedback on the manuscript. We are much grateful to Sven Schenk for his invaluable advice about the treatments with methylfarnesoate. We thank Florian Raible and Alexander Stockinger for their valuable comments on the manuscript. We thank Haley Flom for her diligent proofreading of the manuscript. Animal facility members are thanked for their help with the worm culture. This work was funded by a collaborative grant jointly attributed by the Agence Nationale de la Recherche (ANR) and the Austrian Science Fund (FWF) to the Vervoort (Institut Jacques Monod Paris), Balavoine (Institut Jacques Monod Paris), and Raible (Max Perutz labs Vienna) teams (grant TELOBLAST no. ANR-16-CE91-0007), Labex ‘Who Am I?’ laboratory of excellence (No. ANR-11-LABX-007), Centre National de la Recherche Scientifique, Université de Paris, Association pour la Recherche sur le Cancer (grant PJA 20191209482), and comité départemental de Paris de la Ligue Nationale Contre le Cancer (grant RS20/75-20).

## Data availability statement

All the data are included in the manuscript, figures, supplementary figures, and the supplementary table.

## Supplementary data

**Supplementary Figure 1.**
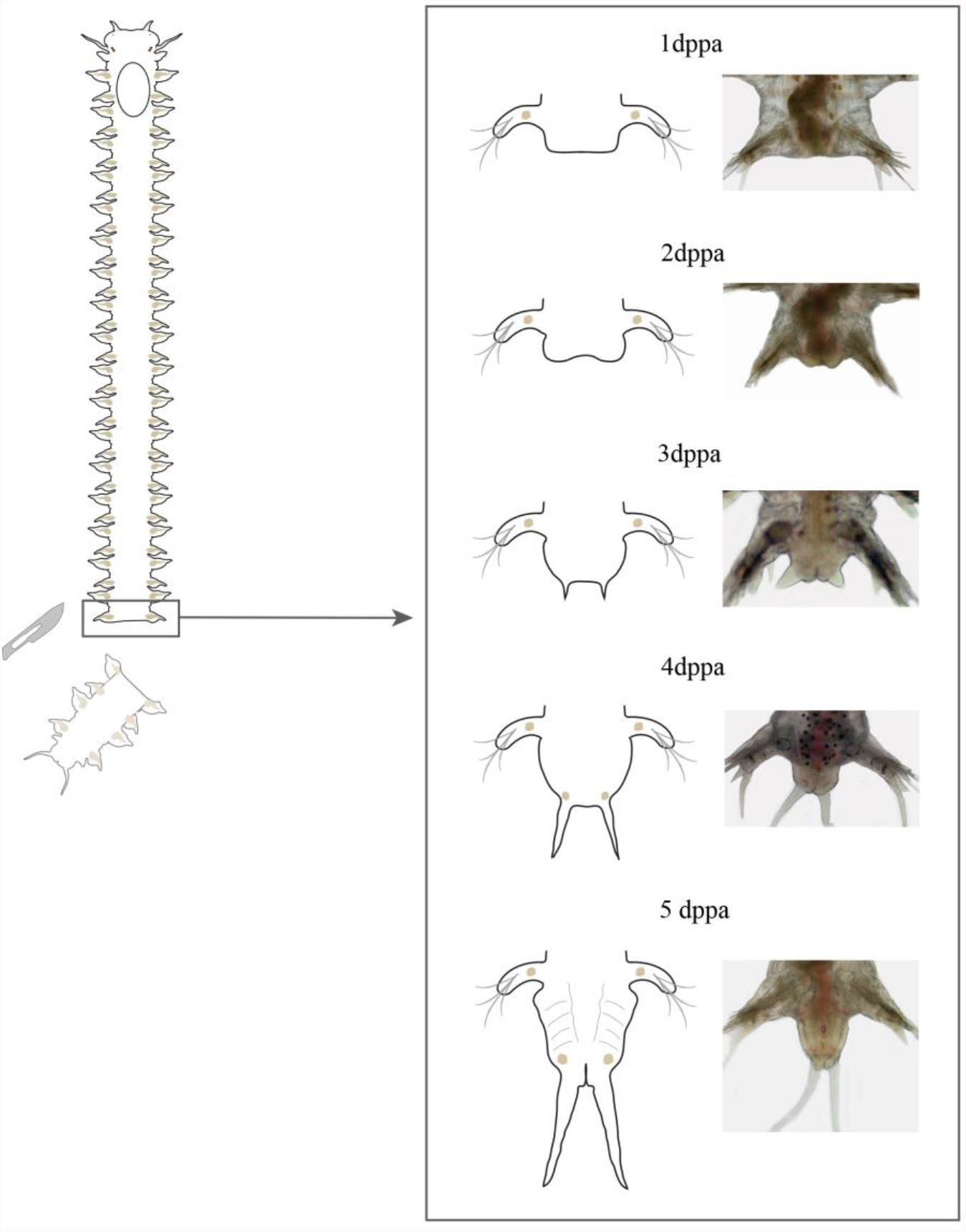
Schematic representation with drawings and bright-field microscopy images of posterior regeneration stages in *P. dumerilii* worms, as defined by Planques et al. 2019. 3-4-months-old juvenile worms are posteriorly amputated (removing around 5 segments, the growth zone and the pygidium). One day post posterior amputation (1 dppa), wound healing is completed; at 2 dppa, the formation of the blastema has started; at 3 dppa, a large blastema with small cirri is present; at 4 dppa, a large blastema containing a differentiated pygidium and long anal cirri is present; at 5 dppa, newly-added segments produced by the regenerated growth zone start to become clearly distinct from the pygidium through the appearance of lateral segmental grooves. At 5 dppa, the regeneration is completed and post regenerative posterior growth starts.

**Supplementary Figure 2.**
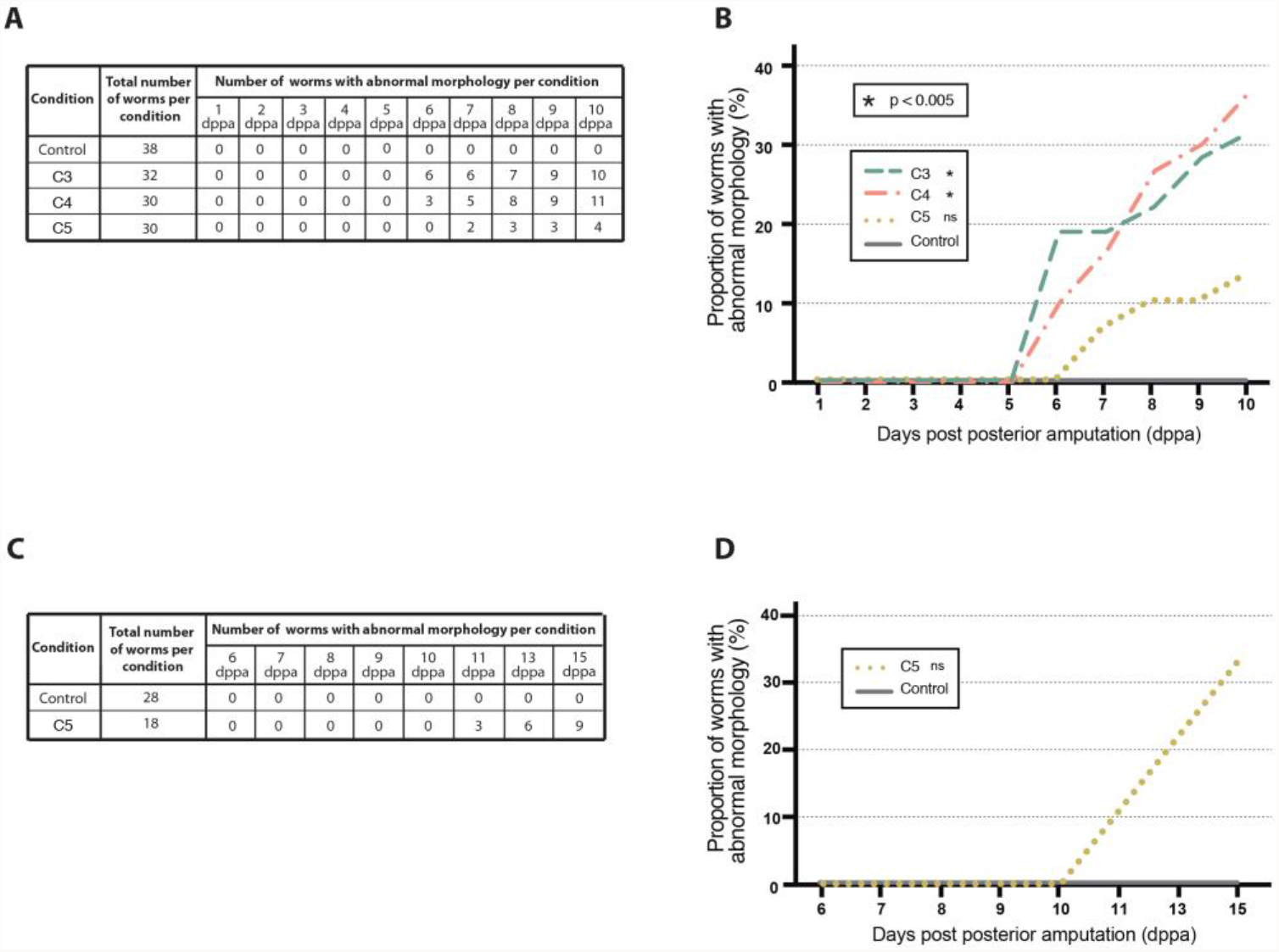
Abnormal morphologies observed during *P. dumerilii* post-regenerative posterior growth after anterior amputation. A) The number of worms showing abnormal morphologies during regeneration (1 to 5 days post posterior amputation, dppa) and post regenerative posterior growth (6 to 10 dppa) for conditions C3 to C5 and control is provided for each scoring day (from 1 to 10 dppa). No worms with abnormal morphology were found in the control condition. The total number of worms per experimental condition is mentioned. Control worms were only amputated posteriorly; for C3, C4 and C5, anterior amputation was made 3, 4 and 5 days after posterior amputation, respectively. See Figure 7 for more details on the experimental designed. B) Graphic representation of the proportion of worms showing abnormal morphologies in condition C3 to C5 from 1 to 10 dppa. Asterisks indicate significant differences in comparison to control, as defined by 2-way ANOVA on repeated measures with Turkey correction. * p < 0.005 (for specific p-values, see Supp. Table 1). C) The number of worms showing abnormal morphologies during post-regenerative posterior growth (6 to 15 dppa) in condition C5 and control is provided for each scoring day. No worms with abnormal morphology were found in the control condition. The total number of worms per experimental condition is mentioned. D) Graphic representation of the proportion of control and C5 worms showing abnormal morphologies during post-regenerative posterior growth, from 6 dppa to 15 dppa. No significant differences were found (for specific p-values, see Supp. Table 1).

**Supplementary Figure 3.**
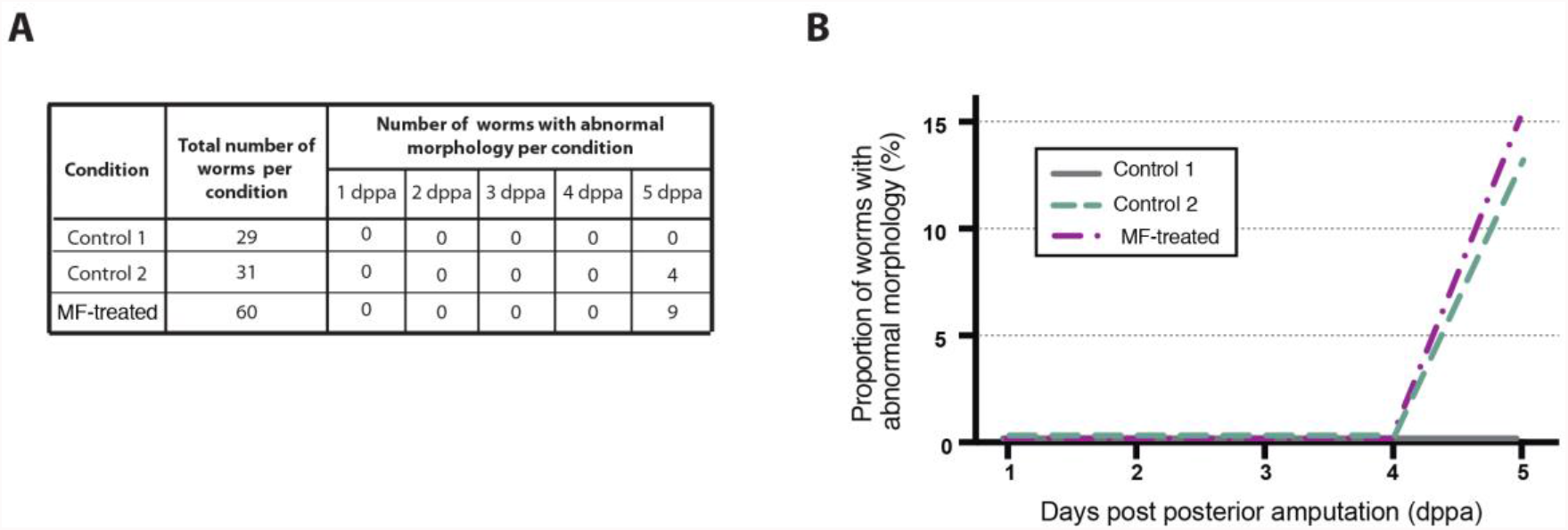
Abnormal morphologies observed during *P. dumerilii* posterior regeneration after anterior amputation and addition of exogenous methylfarnesoate (MF). A) The number of worms showing abnormal morphologies during posterior regeneration (1 to 5 days post posterior amputation, dppa) after anterior amputation and addition of MF (and controls) is provided for each scoring day. Control 1: posterior amputation, no anterior amputation, no MF treatment; Control 2: posterior amputation, anterior amputation, no MF treatment; MF-treated: posterior amputation, anterior amputation, MF treatment (100nM in sea water, renewed every day). No worms with abnormal morphology were found in the control condition. The total number of worms per experimental condition is mentioned. See Figure 9 for more details on the experimental design. B) Graphic representation of the proportion of worms showing abnormal morphologies during posterior regeneration for Control 1, Control 2 and MF-treated conditions from 1 to 5 dppa. No significant differences were found (for specific p-values, see Supp. Table 1).

**Supplementary Table 1**. Results of the 2-way ANOVA statistical analyses performed for each experiment, with the specific correction used for each of them, detailed p-values and significant differences found for the multiple comparisons.

